# Molecular insights into intrinsic transducer-coupling bias in the CXCR4-CXCR7 system

**DOI:** 10.1101/2022.06.06.494935

**Authors:** Parishmita Sarma, Hye-Jin Yoon, Carlo Marion C. Carino, Deeksha S, Ramanuj Banerjee, Yaejin Yun, Jeongsek Ji, Kyungjin Min, Shubhi Pandey, Hemlata Dwivedi-Agnihotri, Xue Rui, Yubo Cao, Kouki Kawakami, Poonam Kumari, Yu-Chih Chen, Kathryn E. Luker, Manish K. Yadav, Ashutosh Ranjan, Madhu Chaturvedi, Jagannath Maharana, Mithu Baidya, Prem N. Yadav, Gary D. Luker, Stéphane A. Laporte, Xin Chen, Asuka Inoue, Hyung Ho Lee, Arun K. Shukla

## Abstract

Chemokine receptors constitute an important subfamily of G protein-coupled receptors (GPCRs), and they are critically involved in a broad range of immune response mechanisms. Ligand promiscuity among these receptors makes them an interesting target to explore novel aspects of biased agonism. Here, we comprehensively characterize two chemokine receptors namely, CXCR4 and CXCR7, which share a common chemokine agonist (CXCL12), in terms of their G-protein coupling, β-arrestin (βarr) recruitment, contribution of GRKs, and ERK1/2 MAP kinase activation. We observe that CXCR7 lacks G-protein coupling while maintaining robust βarr recruitment with a major contribution of GRK5/6. On the other hand, CXCR4 displays robust G-protein activation as expected, however, it exhibits significantly reduced βarr-coupling compared to CXCR7 in response to their shared natural agonist, CXCL12. These two receptors induce distinct βarr conformations even when activated by the same agonist, and CXCR7, unlike CXCR4, fails to activate ERK1/2 MAP kinase. We further determine the crystal structure of βarr2 in complex with a carboxyl-terminal phosphopeptide derived from CXCR7, which reveals a smaller interdomain rotation than observed previously for activated βarrs. Importantly, structure-guided cellular experiments reveal a key contribution of a single phosphorylation site in CXCR7 on βarr recruitment and endosomal trafficking. Taken together, our study provides molecular insights into intrinsic bias encoded in the CXCR4-CXCR7 system, and it has broad implications for therapeutically important framework of biased agonism.

## Introduction

Chemokines are small secreted proteins that typically exert their actions via chemokine receptors belonging to the large superfamily of G protein-coupled receptors (GPCRs), also known as seven transmembrane receptors (7TMRs) (Griffith et al., 2014; Hughes and Nibbs, 2018). Chemokines and chemokine receptors contribute to a diverse array of physiological processes, especially in various aspects of immune response activation and regulation (Griffith et al., 2014; Proudfoot, 2002). A peculiar aspect in the chemokine-chemokine receptor system is ligand promiscuity where not only a single chemokine can bind to, and activate multiple chemokine receptors, but a given receptor can also be activated by several different chemokines (Steen et al., 2014; Zlotnik et al., 2006). Chemokine receptors typically couple to, and signal through, heterotrimeric G-proteins and β-arrestins (βarrs), as expected for prototypical GPCRs (Zhao et al., 2019). Interestingly however, there are several examples of chemokine receptors that exhibit a significant deviation from this paradigm, especially with respect to their transducer-coupling patterns and downstream signaling responses (Cancellieri et al., 2013; Graham et al., 2012; Nibbs and Graham, 2013; Pandey et al., 2021; Ulvmar et al., 2011).

The CXC chemokine receptor subtype 4 (CXCR4) and subtype 7 (CXCR7) constitute an interesting pair as they both recognize a common natural chemokine agonist, referred to as CXCL12, also known as the stromal cell-derived factor 1 (SDF1). These two receptors are involved in various aspects of cancer onset and progression, cardiac disorders and autoimmune diseases (Shi et al., 2020). Interestingly, CXCR7, but not CXCR4, also recognizes another chemokine referred to as CXCL11 (Naumann et al., 2010). CXCR4 is widely considered a prototypical GPCR with coupling to Gαi subfamily of G-proteins as measured in terms of inhibition of cAMP, and it also recruits βarrs upon agonist-stimulation (Luo et al., 2017). On the other hand, stimulation of CXCR7 by CXCL12 or CXCL11 fails to elicit any measurable Gαi activation, although there are reports that suggest its ability to recruit βarrs (Nguyen et al., 2020; Rajagopal et al., 2010). Interestingly, a small molecule ligand known as VUF11207 has also been reported to promote CXCR7-βarr interaction (Wijtmans et al., 2012), although its interaction with other CXCRs and complete transducer-coupling profile has not been evaluated thus far. Therefore, the CXCR4-CXCR7 pair represents an intriguing system to probe the molecular and structural details of intrinsic transducer-coupling bias.

Here, we present a comprehensive investigation of agonist-induced G-protein coupling, βarr recruitment and conformational signatures, ERK1/2 MAP kinase activation, and structural basis of βarr activation by CXCR7. Our study highlights previously unanticipated intrinsic bias encoded in the CXCR4-CXCR7 system, and provides molecular and structural mechanisms for their functional divergence. These findings not only offer important insights to better understand biased agonism at 7TMRs but also present an experimental framework that may guide analogous exploration of other chemokine receptors.

## Results

### Agonist-induced G-protein-activation and second messenger response

CXCR7, unlike CXCR4, is considered to lack G-protein coupling in response to CXCL12 stimulation, although the experimental evidence is limited primarily to the lack of canonical Gαi-activation (Rajagopal et al., 2010) (**Figure 1A**). Therefore, we first set out to comprehensively probe the CXCL12-induced G-protein activation using a previously described NanoBiT-based heterotrimer dissociation assay (Inoue et al., 2019) for CXCR4 and CXCR7. In this assay, agonist-stimulated G-protein activation is measured as a decrease in luminescence signal arising from dissociation of the NanoBiT-engineered heterotrimer consisting of the Gα-LgBiT, SmBiT-Gγ2 and untagged Gβ1 subunits. We observed that CXCR4 robustly activates G-proteins of the Gαi subfamily but CXCR7 remains silent in this assay not only for Gαi but other subtypes as well (**Figure 1B**). As mentioned earlier, a small molecule ligand has also been described for CXCR7 although its characterization remains limited to binding studies and βarr recruitment in a BRET assay (**Figure 1C**). We therefore decided to also test VUF11207 in the NanoBiT-based heterotrimer dissociation assay to probe its ability to activate G-proteins, if any. We observed that like CXCL12, VUF11207 also fails to elicit any measurable G-protein activation from CXCR7 (**Figure 1D**). Moreover, VUF11207 also did not promote any G-protein activation for CXCR4, suggesting its selectivity for CXCR7 (**Figure 1D**). We also tested VUF11207 in second messenger assays based on cAMP response and calcium release, and did not observe a measurable response for CXCR7, which further confirms the inability of CXCR7 to activate G-proteins (**Figure 1E-G**). In these assays, surface expression of both the receptors was optimized to be at comparable levels as measured using either a flow-cytometry based assay (**Figure S1A**) or whole cell-based surface ELISA assay (**Figure S1B-C**). Taken together, these data establish the inherent inability of G-protein activation by CXCR7 upon stimulation by CXCL12 or VUF11207.

**Figure 1.**
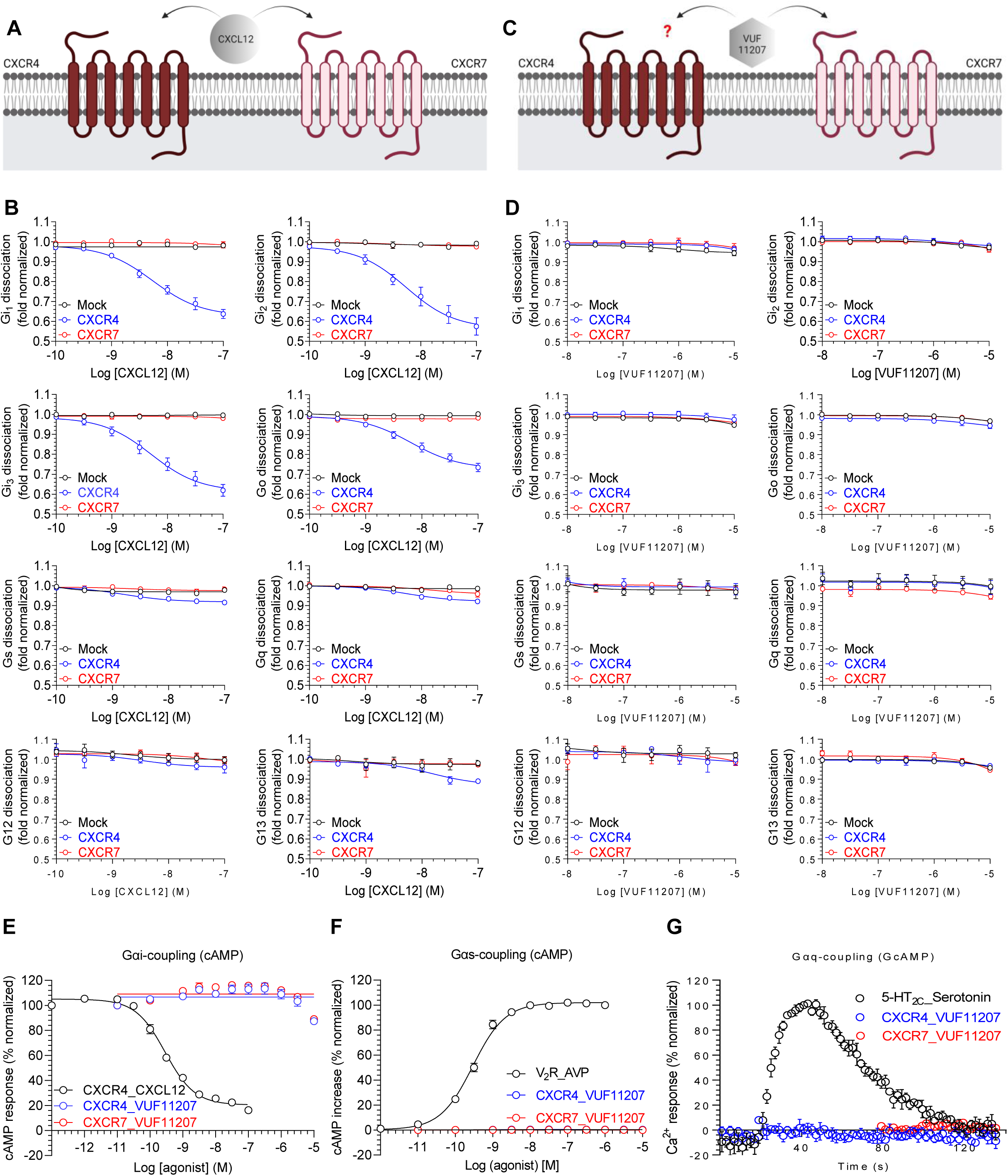
Lack of G-protein coupling to CXCR7. **A.** CXCL12, a CXC type chemokine, is a common agonist for both, CXCR4 and CXCR7. **B.** NanoBiT-based assay for CXCL12-induced dissociation of heterotrimeric G-proteins for CXCR4 and CXCR7 (mean±SEM; n=4-5). Mock represents empty vector transfected cells as a negative control. **C.** VUF11207 is a small molecule agonist for CXCR7 but its efficacy for CXCR4, if any, is not known. **D.** NanoBiT-based assay for VUF11207-induced dissociation of heterotrimeric G-proteins for CXCR4 and CXCR7 (mean±SEM; n=3). **E.** Agonist-induced decrease in forskolin-induced cAMP level measured using the GloSensor assay for the indicated receptor-ligand combinations as a readout of Gαi-activation (mean±SEM; n=4; normalized with starting value for CXCL12-CXCR4 combination as 100%). **F.** Agonist-induced increase in cAMP level measured using the GloSensor assay for the indicated receptor-ligand combinations as a readout of Gαs-activation (mean±SEM; n=3; normalized with maximal signal for V_2_R as 100%) ). V_2_R (vasopressin receptor subtype 2) is used as a positive control. **G.** Agonist-induced increase in Ca^2+^ level measured using the GcAMP sensor for the indicated receptor-ligand combinations as a readout of Gαq-activation (mean±SEM; n=4; normalized with maximal signal for serotonin as 100%). 5-HT_2C_ receptor is used as a positive control.

### Agonist-induced βarr recruitment

Although previous studies have shown βarr coupling to CXCR4 (Luo et al., 2017) and CXCR7 (Nguyen et al., 2020), a comprehensive side-by-side analysis of recruitment of both βarr isoforms i.e. βarr1 and 2, to both receptors has not been described thus far. Therefore, we used two different assays to measure recruitment of βarr1 and 2 to both these receptors in response to CXCL12 and VUF11207. First, we measured CXCL12-induced βarr2 recruitment in a previously described PRESTO-Tango assay (Kroeze et al., 2015), where the receptors are engineered to contain the carboxyl-terminus of vasopressin subtype 2 receptor (i.e. CXCR4-V_2_ and CXCR7-V_2_). While we observed a robust response for CXCR7 in a ligand dose-dependent manner, the basal luminescence signal in CXCR4-expressing cells was high and it did not change much in response to CXCL12 (**Figure 2A**). Considering that PRESTO Tango constructs use a chimeric receptor, we also generated new Tango assay constructs for CXCR4 and CXCR7 without the V_2_R carboxyl-terminus fusion, and measured βarr2 recruitment in response to CXCL12. We observed a robust agonist-induced response for CXCR7, however, CXCR4 displayed a significantly lower E_max_ for βarr2 recruitment compared to CXCR7 (**Figure 2B**). On the other hand, VUF11207 effectively promoted βarr2 recruitment for CXCR7 but not for CXCR4 further suggesting its specificity at CXCR7 (**Figure 2C-D**).

**Figure 2.**
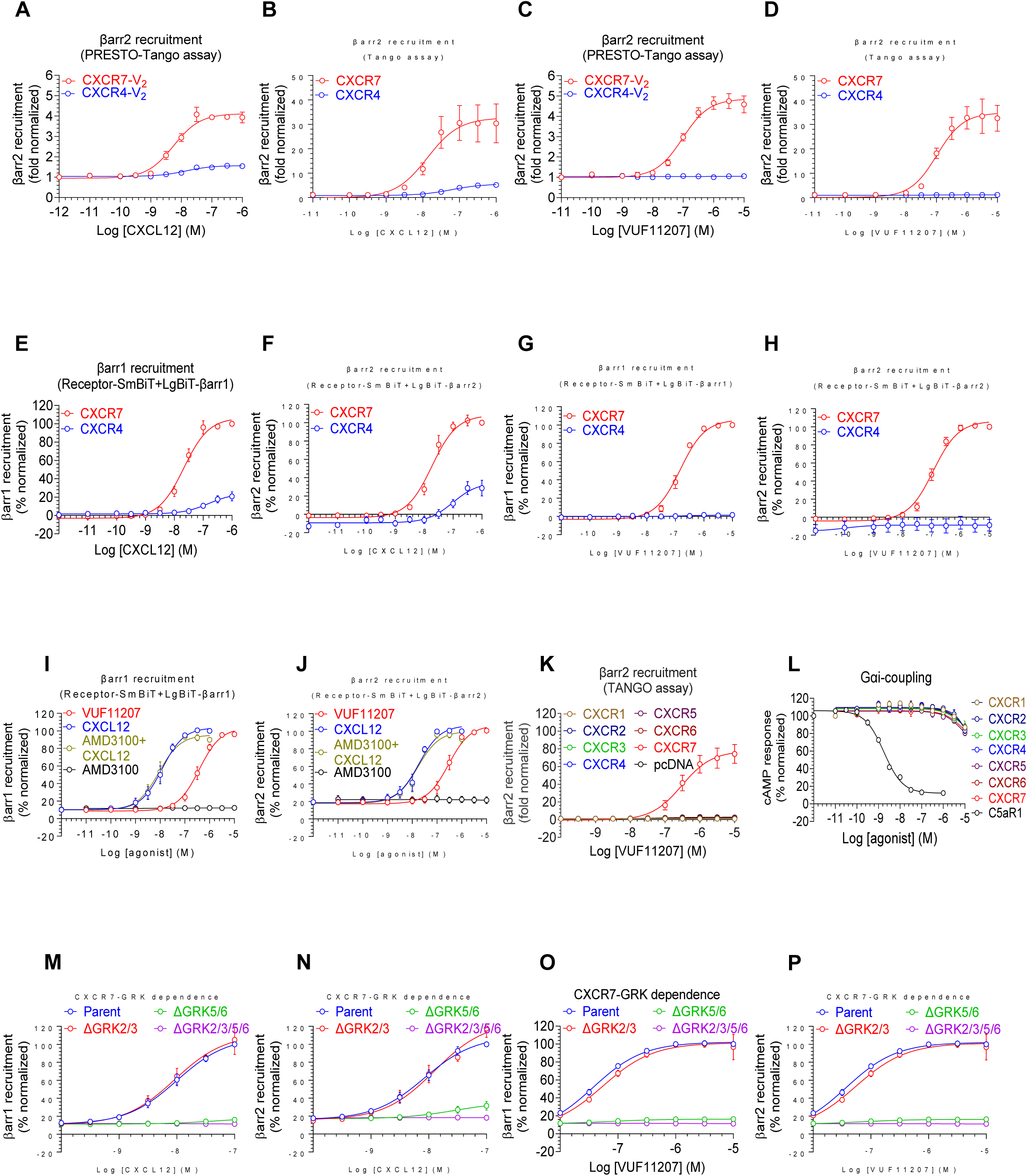
β-arrestin recruitment to CXCR7. **A-B.** CXCL12-induced βarr2 recruitment to CXCR4 and CXCR7 in PRESTO-Tango and Tango assays, respectively (mean±SEM; n=3; normalized with the luminescence signal at minimal ligand dose as 1). The PRESTO-Tango assay uses a chimeric receptor construct with the carboxyl-terminus of V_2_R while the Tango assay uses native receptors. **C-D.** VUF11207-induced βarr2 recruitment to CXCR4 and CXCR7 in PRESTO-Tango and Tango assays, respectively (mean±SEM; n=3; normalized with the luminescence signal at minimal ligand dose as 1). **E-F.** CXCL12-induced βarr1/2 recruitment to CXCR4 and CXCR7 in NanoBiT assay (Receptor-SmBiT+LgBiT-βarr1/2), respectively (mean±SEM; n=4; normalized with luminescence signal for CXCR7 at maximal ligand dose as 100%). **G-H.** VUF11207-induced βarr1/2 recruitment to CXCR4 and CXCR7 in NanoBiT assay (Receptor-SmBiT+LgBiT-βarr1/2), respectively (mean±SEM; n=4; normalized with the maximal ligand dose luminescence signal for CXCR7 as 100%). **I-J.** A side-by-side comparison of CXCL12- vs. VUF11207-induced βarr1 and 2 recruitment to CXCR7, respectively (mean±SEM; n=4; normalized with the maximal ligand dose luminescence signal for CXCR7 as 100%) (pEC_50_ for βarr1: VUF 11207, 6.42±0.05, CXCL12, 7.72±0.07; βarr2, VUF11207, 6.53±0.08, CXCL12: 7.7±0.08). A CXCR4-specific antagonist AMD3100 is used either alone, or as pre-treatment to CXCL12, as a negative control and to rule out the possibility of any contribution from endogenous CXCR4. **K.** VUF11207-induced βarr2 recruitment for all the CXC chemokine receptors (CXCR1-7) in the Tango assay (mean±SEM; n=5; normalized with the luminescence signal at minimal ligand dose as 1. **L.** VUF11207-induced Gαi-coupling for all the CXC chemokine receptors (CXCR1-7) in the GloSensor assay (mean±SEM; n=3; normalized with the starting value for each receptor as 100%). **M-N.** CXCL12-induced βarr1 and 2 recruitment to CXCR7 in GRK knock-out cells using the NanoBiT assay (Receptor-SmBiT+LgBiT-βarr1/2), respectively (mean±SEM; n=4-5; normalized with the maximal ligand dose luminescence signal for CXCR7 at parent condition as 100%). **O-P.** VUF11207-induced βarr1 and 2 recruitment to CXCR7 in GRK knock-out cells using the NanoBiT assay (Receptor-SmBiT+ LgBiT-βarr1/2), respectively (mean±SEM; n=5; normalized with the maximal ligand dose luminescence signal for CXCR7 at parent condition as 100%).

We found the weaker βarr2 recruitment response for CXCR4 compared to CXCR7 in the Tango assay intriguing, and therefore, we generated NanoBiT assay constructs for CXCR4 and CXCR7, which enable us to measure βarr response in real time, to probe this further. In this assay, the receptors were tagged with the SmBiT at their carboxyl-terminus and they were co-expressed with LgBiT-tagged βarr1 or βarr2 followed by the measurement of agonist-induced change in the luminescence signal. Similar to Tango assay data, we observed a stronger recruitment in terms of E_max_ of βarr1 and 2 for CXCR7 compared to CXCR4 upon stimulation with CXCL12 (**Figure 2E-F**) while VUF11207 selectively promoted βarr1 and 2 recruitment to CXCR7 but not to CXCR4 (**Figure 2G-H**). These data not only suggest the selectivity of VUF11207 for CXCR7 but more importantly, also underscore a relatively lower propensity of βarr-coupling to CXCR4 compared to CXCR7. In addition, a closer analysis of βarr recruitment to CXCR7 in the NanoBiT assay suggested a difference in the EC_50_ between CXCL12 and VUF11207. Therefore, we compared these two ligands side-by-side and confirmed that CXCL12 is more potent for βarr recruitment over VUF11207, although their E_max_ values are comparable (**Figure 2I-J**). In all these assays, CXCR4 and CXCR7 were expressed at comparable levels as measured by whole cell-based surface ELISA assay (**Figure S1D-G**).

### Selectivity profiling of VUF11207 on CXCRs

Inspired by the selectivity of VUF11207 for CXCR7 over CXCR4, we decided to test it on other CXCRs as well. There are seven CXCRs (CXCR1-CXCR7) in the human genome, and we measured VUF11207 response for all of them in parallel using the GloSensor assay for Gαi-coupling and PRESTO-Tango/Tango assay for βarr2 coupling. We observed that VUF11207 was able to induce βarr2 recruitment only for CXCR7 and no other CXCRs (**Figure 2K** **and Figure S2C**). Moreover, VUF11207 was completely silent for every CXCR tested in the GloSensor-based cAMP assay as a readout of Gαi-coupling (**Figure 2L**). We observed some variations in the relative receptor expression in these assays, although they all expressed at levels that are higher than the mock-transfected cells (**Figure S2A-B and S2D**). Taken together, these findings establish VUF11207 as a CXCR7 selective agonist and therefore, provides a pre-validated tool compound to probe the structure and function of CXCR7 in the future studies.

CXCR7 promotes cellular migration upon activation by CXCL12, and therefore, we also measured if VUF11207 promotes migration in cells expressing CXCR7. We measured the migration of MDA-MB-231 breast cancer cells that were stably transfected with CXCR7 using a 2D microfluidic device as described previously (Song et al., 2009). We observed that stimulation of these cells with VUF11207 resulted in efficient migration as compared to vehicle-treated cells, and that VUF11207 response was comparable to that of CXCL12 (**Figure S2E**). Simultaneous addition of CXCL12 and VUF11207 did not result in any synergistic effect on migration of these cells (**Figure S2E**).

### Contribution of GRKs in βarr recruitment

As receptor phosphorylation is a key determinant of βarr recruitment, we next tested the contribution of different GRKs in agonist-induced βarr recruitment to CXCR7 using recently described GRK knock-out cell lines (Kawakami et al., 2022). We observed that knock-out of GRK5/6 nearly ablates CXCR7-βarr interaction in response to both agonists, CXCL12 and VUF11207 (**Figure 2M-P**). On the other hand, knock-out of GRK2/3 did not influence βarr recruitment to CXCR7 (**Figure 2M-P**). This observation converges with the lack of G-protein-coupling to CXCR7 because GRK2/3 translocation to the plasma membrane and activation and trafficking has previously been shown to require Gβγ release (Pitcher et al., 1992). We also assessed agonist-induced CXCR7-βarr interaction in presence of pertussis toxin (PTX), however, the pattern of recruitment in response to either CXCL12 or VUF11207 did not change significantly (**Figure S3A-D**). This observation rules out activation-independent contribution of Gαi on βarr recruitment, and therefore, establishes CXCR7 as a model βarr-coupled receptor system to further investigate the functional contribution of βarrs.

### Agonist-induced ERK1/2 activation and conformations of βarr2 for CXCR4 vs. CXCR7

In order to test if βarr recruitment results in ERK1/2 MAP kinase activation, we measured agonist-induced ERK1/2 phosphorylation for CXCR4 and CXCR7 in response to CXCL12 in transfected HEK293 cells. Although we observed robust ERK1/2 phosphorylation in CXCR4- and CXCR7-expressing cells, there was a similar response in mock-transfected cells as well (**Figure 3A-B**). Moreover, ERK1/2 phosphorylation was effectively blocked by the pre-treatment of AMD3100 (CXCR4 antagonist) (**Figure 3C-D** **and S4A-B**). Furthermore, CXCL12-induced ERK1/2 phosphorylation in CXCR4-expressing cells was also completely abolished by pre-treatment with pertussis toxin (PTX) suggesting a major dependence on Gαi activation (**Figure S3E-F**). Thus, CXCL12-induced ERK1/2 phosphorylation likely arises from the endogenous CXCR4 expressed in HEK293 cells as suggested previously(Nguyen et al., 2020). Interestingly, VUF11207 failed to elicit any measurable ERK1/2 activation at either CXCR4 or CXCR7 (**Figure 3E-F**). In these experiments, the receptors were expressed at significantly higher level than mock-transfected cells (**Figure S4C-D**). While the lack of VUF11207-induced ERK1/2 response at CXCR4 is expected, its inability to activate ERK1/2 at CXCR7, in spite of robust βarr recruitment, is intriguing. Taken together, these data reveal that CXCR7 activation, either by CXCL12 or VUF11207, does not initiate ERK1/2 MAP kinase activation despite robust βarr recruitment. These findings also underscore the notion that βarr recruitment does not always translate to ERK1/2 activation and it may depend on specific conformation adopted by βarrs in complex with 7TMRs.

**Figure 3.**
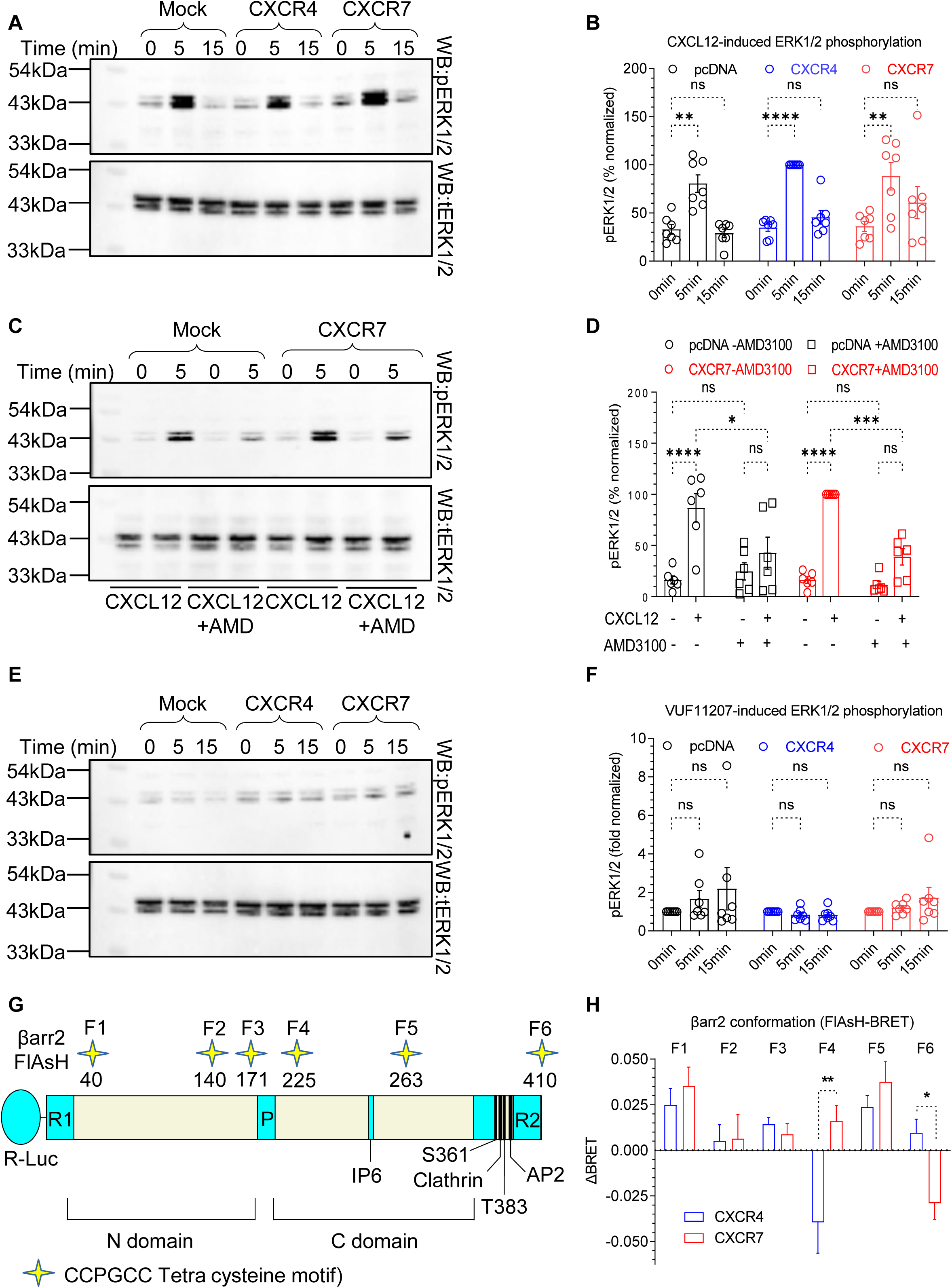
Agonist-induced ERK1/2 phosphorylation and βarr2 conformation for CXCR4 and CXCR7. **A.** HEK293 cells expressing CXCR4 and CXCR7 exhibit a robust signal for ERK1/2 phosphorylation as measured by Western blotting, however, mock-transfected cells also show similar levels of ERK1/2 phosphorylation suggesting that the response is driven primarily via the endogenous CXCR4. **B.** Densitometry-based quantification of ERK1/2 phosphorylation (mean±SEM; n=7, normalized with the 5 min signal for CXCR4 as 100%, Two-way ANOVA, Tukey’s multiple comparison test, **p<0.01, **** p<0.0001, ns= non-significant ). **C.** CXCL12-induced ERK1/2 phosphorylation in CXCR7 expressing cells is blocked by pre-treatment with AMD3100 (10 μM, 30 min). **D.** Densitometry-based quantification of ERK1/2 phosphorylation (mean±SEM; n=6, normalized with CXCL12-induced signal for CXCR7 as 100%, Two-way ANOVA, Tukey’s multiple comparison test, *p<0.05 ***p<0.001, **** p<0.0001, ns= non-significant ). **E.** VUF11207-induced ERK1/2 phosphorylation in HEK293 cells expressing CXCR4 and CXCR7 as measured by Western blotting. **F.** Densitometry-based quantification of ERK1/2 phosphorylation data presented in panel E (mean±SEM; n=6, normalized with respect to the 0 min signal for each condition treated as 1, Two-way ANOVA, Tukey’s multiple comparison test, ns= non-significant ). **G.** A schematic representation of intramolecular BRET-based sensors of βarr2 conformation where the N-terminus of βarr2 harbors R-Luc (Renilla luciferase) as the BRET donor, and the FlAsH motif are engineered at indicated positions in βarr2 as BRET acceptor. **H.** CXCL12-induced BRET signal measured in HEK293 cells expressing the indicated receptor and sensor constructs (mean±SEM, n=6; Two-way ANOVA, Sidak’s multiple comparison test; *p<0.05, **p<0.01 ).

Prompted by the lack of ERK1/2 phosphorylation despite robust βarr recruitment, we decided to probe the conformation of βarr2 upon its interaction with CXCR4 and CXCR7. We used a BRET-based approach where tetra-cysteine FlAsH motifs are engineered at six distinct sites in βarr2 as BRET acceptor while the Renilla luciferase (R-Luc) is engineered at the N-terminus (Lee et al., 2016; Nuber et al., 2016) (**Figure 3G**). A change in BRET signal therefore reports conformational changes in βarr2, and a side-by-side comparison of BRET signal for these six different sensors offers readout of conformational changes in βarr2 imparted by its interaction with a receptor. Interestingly, we observed that the changes in BRET signal for several of these sensors were significantly different between CXCR4 and CXCR7 in response to CXCL12 (**Figure 3H**). For example, in case of sensor F4 where the FlAsH label is localized between β-strand 14 and 15, there was a decrease in BRET signal for CXCR4 but an increase for CXCR7. Similarly, in case of sensor F6 where the FlAsH label is localized at position 410 in the carboxyl-terminus, there was an increase in BRET signal for CXCR4 but a decrease for CXCR7. In other sensors, the pattern of BRET change was similar between the two receptors. Taken together, these data suggest that βarr2 adopts distinct conformations upon its recruitment to CXCR4 vs. CXCR7 in response to CXCL12.

### Identification of the key phosphorylation-site cluster in CXCR7

As receptor phosphorylation is a key determinant of βarr binding, we analyzed the carboxyl-terminus of CXCR7 and identified two distinct phosphorylation clusters, each containing three potential phosphorylation sites (**Figure 4A**). These two clusters, referred to as cluster 1 and 2 from here onwards, harbor PXXPXXP and PXPXXP type pattern of phosphorylation sites, respectively, where P represents a Serine or Threonine. We generated two different CXCR7 constructs by mutating the phosphorylation sites in these clusters, and monitored agonist-induced trafficking of βarr2 compared to the wild-type CXCR7. We observed that CXCR7 mutant lacking cluster 1 induces a βarr2 trafficking pattern that is nearly identical to wild type receptor, however, mutation of cluster 2 completely abrogates βarr2 trafficking in response to either CXCL12 or VUF11207 (**Figure 4B-C**). These data establish cluster 2 as the major determinant of βarr2 interaction and trafficking for CXCR7, and prompted us to investigate the structural mechanism at high-resolution.

**Figure 4.**
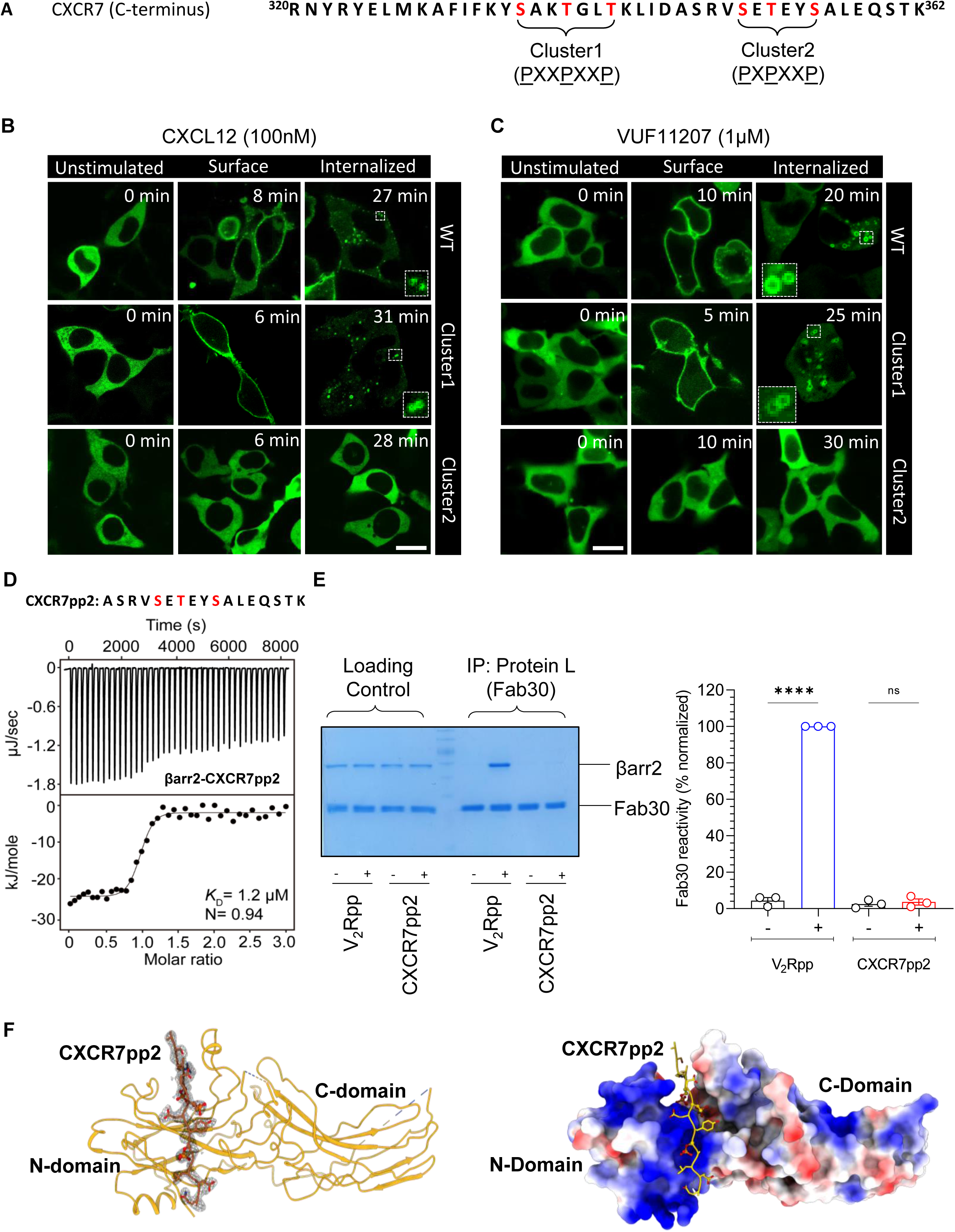
Key phosphorylation site cluster in CXCR7 and crystal structure of βarr2. **A.** The carboxyl-terminus of CXCR7 harbors two potential phosphorylation site clusters indicated as cluster 1 and 2 with PXXPXXP and PXPXXP type phosphorylation codes (P is Ser/Thr; X is any other amino acid). **B-C.** CXCL12- and VUF11207-induced trafficking of βarr2 as monitored using confocal microscopy in HEK293 cells expressing the indicated receptor constructs (WT=wild-type CXCR7; Cluster1=CXCR7 with S^335^A/T^338^A/T^341^A mutations; Cluster2= CXCR7 with S^350^A/T^352^A/T^355^A mutations). Representative images from three independent experiments are shown. Scale bar is 10μm. **D.** The cluster2 phosphopeptide (CXCR7pp2) binds βarr2 with high affinity as measured by isothermal calorimetry (ITC). Purified βarr2 was incubated with increasing concentration of the peptide and their binding parameters were calculated based on the dose response curve. **E.** Fab30 fails to recognize CXCR7pp2-bound βarr2 although it robustly pulls down V_2_Rpp-bound βarr2 as measured using co-immunoprecipitation (co-IP) assay. Purified βarr2 was incubated with saturating concentration of the indicated phosphopeptides followed by addition of 1.5-fold molar excess of Fab30 and co-IP using Protein-L agarose. A representative image and densitometry-based quantification is presented here (mean±SEM; n=3, normalized with V_2_Rpp signal as 100%). Data are analyzed using One-way ANOVA with Sidak’s multiple comparison test(****p<0.0001, ns= non-significant). **F.** Ribbon diagram of the CXCR7pp2-bound βarr2 crystal structure showing the overall peptide docking interface on the N-domain. The right panel shows the surface representation with overall charge distribution.

### Crystal structure of CXCR7pp2-βarr2 complex

We synthesized a phosphopeptide corresponding to the cluster 2, referred to as CXCR7pp2, and measured its interaction with C-terminally truncated and pre-activated βarr2 construct (Δ357-410) using isothermal titration calorimetry (ITC). We observed a monophasic binding of CXCR7pp2 with an affinity of approximately 1.2 µM (**Figure 4D**). In order to facilitate the crystallization of CXCR7pp2-βarr2 complex, we attempted to use a previously described antibody fragment (Fab30) that has been used to obtain a high-resolution structure of V_2_Rpp-βarr1 complex (Shukla et al., 2013) and also several GPCR-βarr complexes (Lee et al., 2020; Staus et al., 2020). Surprisingly however, we observed that Fab30 does not recognize CXCR7pp2-bound βarr2 conformation as measured in a co-immunoprecipitation (co-IP) assay while it robustly recognized V_2_Rpp-βarr2 complex as reported previously(Ghosh et al., 2019) (**Figure 4E**). This finding further agrees with the BRET data (**Figure 3G-H**) in terms of a distinct βarr2 conformation induced by CXCR7 compared to a prototypical GPCR. Therefore, we attempted to crystallize the CXCR7pp2-βarr2 complex using a C-terminally truncated and pre-activated βarr2 construct (Δ357-410), and successfully obtained a crystal structure at 2.8 Å resolution (**Figure 4F**).

The analysis of intensity statistics of the X-ray diffraction data with Phenix Xtriage suggested possible twinning, related by pseudo-merohedral twin operator -h, -k, l. In the crystal structure, we observed six molecules in the asymmetric unit, which are labeled as A-F, and the corresponding phosphopeptide chains are labeled as U-Z (**Figure S5A**). For structural analysis and interpretation, we focused on chain A and D of βarr2, and the corresponding chain U and X of the CXCR7pp2. Compared to the previously determined crystal structure of βarr2 in a basal state(Zhan et al., 2011), CXCR7pp2-bound βarr2 displays approximately 8° of inter-domain rotation between the N- and the C-domain (**Figure 5A**). This is significantly smaller than the inter-domain rotation observed not only in the V_2_Rpp-βarr1 complex(He et al., 2021; Shukla et al., 2013) but also in β_1_AR-βarr1, NTS_1_R-βarr1(Huang et al., 2020) and IP_6_-bound βarr2(Chen et al., 2017) (**Figure 5B**). For measuring the inter-domain rotation in various structures, we superimposed the N-domain of βarrs in different structures and fixed their position relative to the N-domain of the corresponding basal state structures (i.e. 1G4M for βarr1 and 3P2D for βarr2). This was followed by rotation measurement across the C-domains taking residue Pro^179^ as the hinge. It is interesting to note that a previous study has suggested a direct correlation between the inter-domain rotation in βarrs with Fab30 reactivity, and an inter-domain rotation of 15° or higher appears to support Fab30 binding (Dwivedi-Agnihotri et al., 2020). In this regard, the lack of Fab30 binding to CXCR7pp2-βarr2 complex (**Figure 4E**) agrees well with the smaller inter-domain rotation observed in the crystal structure.

**Figure 5.**
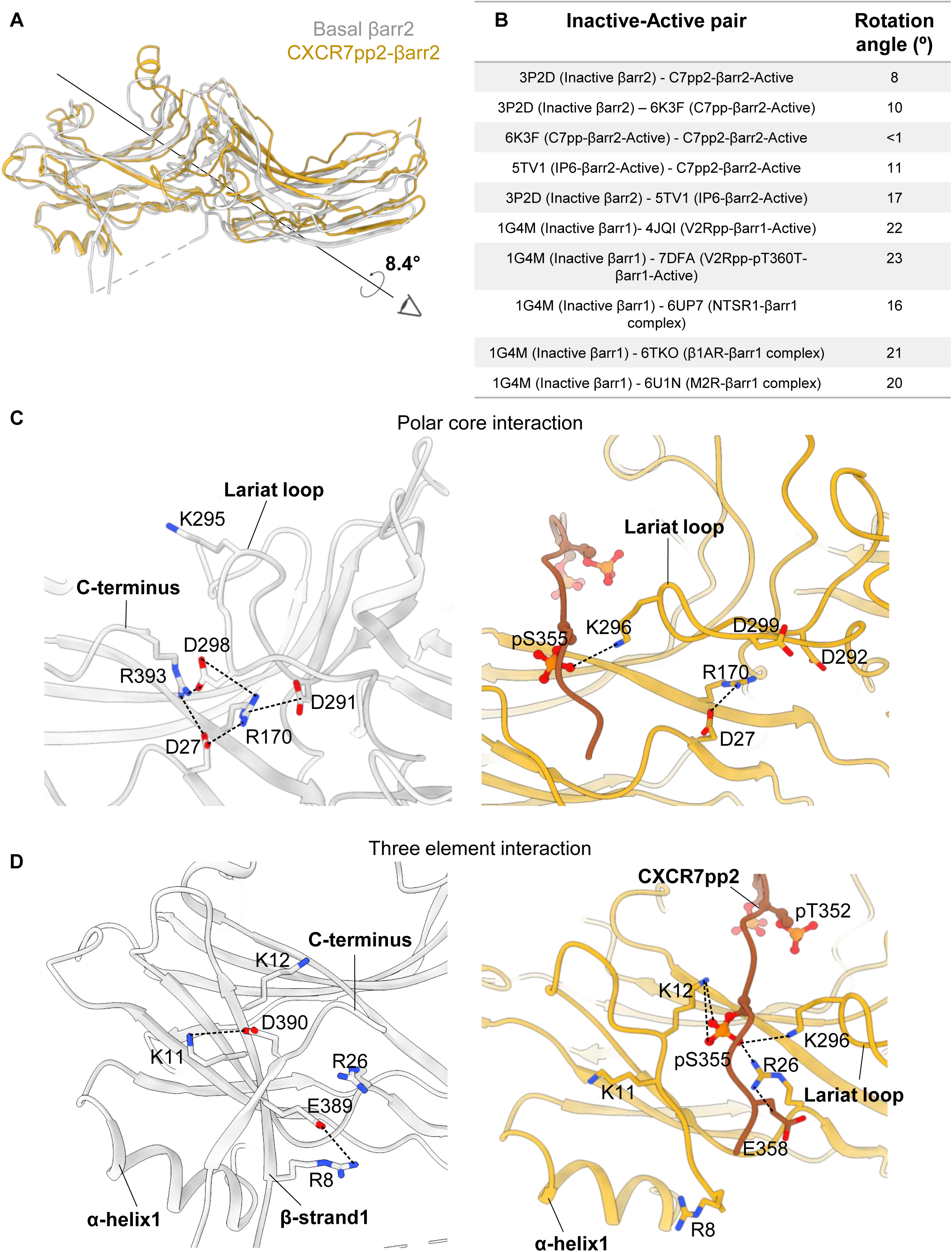
Activation-induced conformational changes in βarr2 as observed in the crystal structure. **A.** Superimposition of CXCR7pp2-bound βarr2 structure with the basal conformation (PDB: 3P2D) reveals an interdomain rotation of about 8.4° calculated using a script in PyMol. **B.** Comparison of the inter-domain rotation in CXCR7pp2-βarr2 crystal structure with previously determined structures of βarrs calculated using PyMol script. **C.** Structural comparison of the polar core interaction between the basal (gray) and CXCR7pp2-bound βarr2 conformations (orange). Binding of CXCR7pp2 disrupts the polar core network by displacing the carboxyl-terminus of βarr2, which results in the loss of R^393^-D^298^ interaction and a significant reorientation of the lariat loop. **D.** Structural comparison of the three element interaction in the N-domain of βarr2 between the basal and CXCR7pp2-bound conformations. CXCR7pp2 displaces the carboxyl-terminus of βarr2, and the three phosphate groups make multiple ionic interactions with the Lys/Arg in βarr2.

One of the hallmarks of the basal conformation of βarrs is a network of interactions referred to as the polar core, and βarr activation leads to the disruption of this network (Zhan et al., 2011). The polar core in βarr2 comprises of charged residues from the N-domain (Asp^27^ and Arg^170^), C-domain (Asp^291^ and Asp^298^), and the carboxyl-terminus (Arg^393^), bringing different parts of βarr2 together i.e. N-domain, C-domain, and the C-terminus (**Figure 5C**). Among these, Arg^393^ is involved in two different salt-bridge interactions with Asp^27^ (N-domain) and Asp^298^ (lariat loop) (**Figure 5C**). The previously determined crystal structure of βarr2 (PDB ID: 3P2D) is of bovine protein while we have used rat βarr2 in our studies. The insertion of Arg^197^ in rat βarr2 results in a shift of the amino acid numbering by one from position 197 onwards. As we used truncated βarr2 for crystallization, it lacks Arg^394^ and therefore, expectedly leads to partial disruption of the polar core. In addition, we also observed that pSer^355^ in CXCR7pp2 engages with Lys^296^ in βarr2 through a salt-bridge, which repositions the lariat loop leading to the disruption of Arg^170^-Asp^299^ and Arg^170^-Asp^292^ interactions involved in the polar-core (**Figure 5C**). However, the Asp^27^-Arg^170^ interaction remains intact even after CXCR7pp2 binding (**Figure 5C**). Another key interaction that restrains βarrs in their basal state is referred to as the three-element interaction consisting of β-strand I and α-helix I in the N-domain and β-strand XX in the carboxyl-terminus of βarrs (**Figure 5D**). Here, Arg^8^ and Lys^11^ on β-strand I form ionic interactions with Glu^389^ and Asp^390^, respectively, which are localized in the carboxyl-terminus of βarr2. In the CXCR7pp2-βarr2 structure, the three-element interaction is partly disrupted due to the lack of carboxyl-terminus, which is also reflected by the reorientation of the side-chains of Arg^8^ and Lys^11^ (**Figure 5D**). In addition, we also observed that Arg^26^ engages in new interactions with pSer^355^ and Glu^358^ in the CXCR7pp2 while Lys^12^ interacts with pSer^355^ (**Figure 5D**).

In order to gain further structural insights into βarr2 activation, we compared the overall conformation of three major loops (finger loop, middle loop and lariat loop) in βarr2 between the CXCR7pp2-bound structure and the basal state (**Figure 6A**). We observed significant reorientation of these loops upon CXCR7pp2-binding. For example, there was an outward movement of the finger loop upon βarr2 activation and the appearance of a helical turn that has also been observed in NTS_1_R-βarr1 structure (Huang et al., 2020) and IP6-bound βarr2 (Chen et al., 2017). The lariat loop moved closer to the N-domain, possibly due to the interaction of Lys^296^ with the pSer^355^ in CXCR7pp2. The middle loop is also shifted significantly in the CXCR7pp2-bound structure compared to the basal state, similar to that observed in other activated βarr structures (Lee et al., 2020; Shukla et al., 2013). Finally, the hydrogen-bond network stabilized by the intramolecular interactions among the finger loop, middle loop, and the C-loop is also broken in CXCR7pp2-βarr2 structure, which allows the significant reorientation of the key loops as discussed above. It is also worth noting that the overall conformation and orientation of these loops matched well among the six molecules in the asymmetric unit, suggesting that the observed conformations were not influenced by crystal packing and crystallographic contacts (**Figure S5B and S6**).

**Figure 6.**
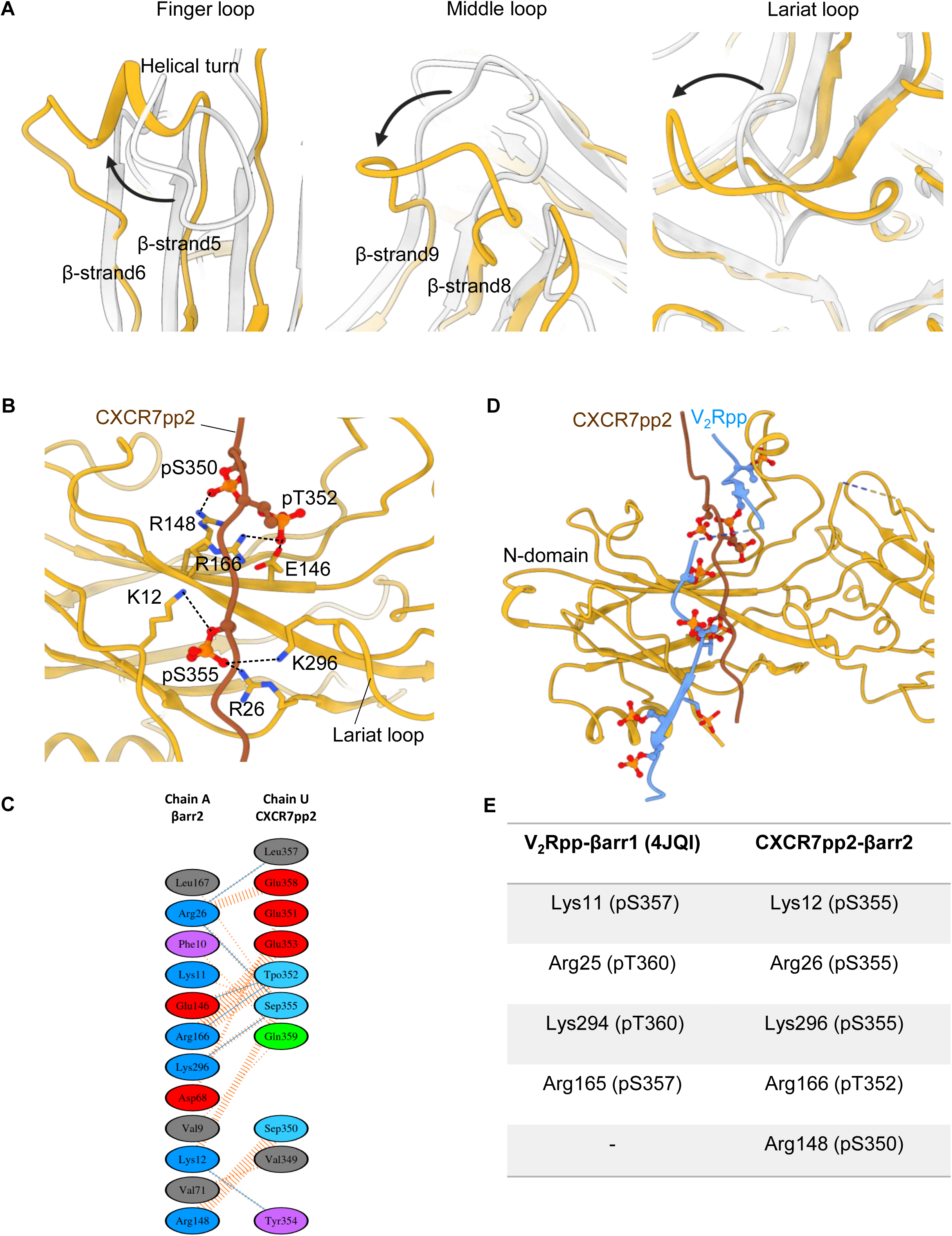
Structural details of activation-induced conformational changes in βarr2 as observed in the crystal structure. **A.** Reorientation of the finger, middle and the lariat loop in βarr2 upon CXCR7pp2 binding. Structures of βarr2 in the basal and CXCR7pp2-bound states were superimposed and the relative positioning of the indicated loops are presented. **B.** A structural snapshot of CXCR7pp2-βarr2 structure depicting the interaction of phosphates with Lys/Arg in βarr2. The dotted lines indicated ionic interactions between the labeled residues. **C.** Overall interaction network of CXCR7pp2 with βarr2 as calculated using PDBSum based on the crystal structure of CXCR7pp2-βarr2 structure. **D.** Structural comparison of overall docking mode of the V_2_Rpp on βarr1 (PDB:4JQI) and CXCR7pp2 on βarr2 suggest a similar binding pocket on the N-domain. **E.** A comparison of the phosphate-interacting residues in CXCR7pp2-βarr2 structure with that of V_2_Rpp-βarr1 structure. Analogous Lys/Arg residues that interact with the phosphates in both the structures are presented.

We next analyzed the interaction pattern of phosphate groups present in CXCR7pp2 with the different amino acids in βarr2. CXCR7pp2 contains three phosphates, which consist of frequently observed phosphorylation pattern (PXPXXP) in the carboxyl-terminus of GPCRs (Zhou et al., 2017). We observed that pSer^350^ forms a salt-bridge with Arg^148^, pThr^352^ with Arg^166^, and pSer^355^ with Lys^12^, Arg^26^ and Lys^296^ (**Figure 6B-C**). A comparison of CXCR7pp2-bound βarr2 with the crystal structure of V_2_Rpp-βarr1 complex reveals several interesting observations. For example, the β sheet stacking of N- and C-terminal parts of the V_2_Rpp is absent in CXCR7pp2 although CXCR7pp2 (17 amino acids) is also significantly smaller than V_2_Rpp (29 amino acids), and it contains lesser number of phosphates (**Figure 6D**). Moreover, the N-terminal part of CXCR7pp2 is away from the finger loop and closer to the β7/β8 loop compared to the V_2_Rpp (**Figure 6D**). Strikingly, the interaction of the phosphate groups in the CXCR7pp2 with Lys^12^, Arg^26^, Arg^166^ and Lys^296^ are analogous to that observed in V_2_Rpp-βarr1 structure, suggesting a potentially conserved role of these residues as phosphate sensors in βarr isoforms (**Figure 6E**).

### Structure-guided identification of a key phosphorylation site in CXCR7

In order to further validate the structural observations, we generated a series of CXCR7 mutants lacking either one or a combination of phosphorylation sites, and tested their ability to recruit βarrs in Tango and NanoBiT assay upon stimulation with CXCL12 and VUF11207 (**Figure 7A-D**). We observed that the mutation of a single phosphorylation site i.e. Thr^352^ resulted in a dramatic decrease in βarr2 recruitment, while the other two sites had relatively modest effect individually (**Figure 7A-D**). A combination of two sites where one was Thr^352^ had an additive effect in terms of attenuation in βarr2 binding, while the mutant lacking all three phosphorylation sites completely lost βarr2 recruitment. In addition to the data presented in (**Figure 7A-D**), where a two saturating doses of agonists were used, we also carried out complete dose response experiments on selected mutants, which further recapitulated the same pattern of βarr2 recruitment (**Figure 7E-H**). In order to test if diminished βarr2 recruitment also translates to reduced functional response, we measured agonist-induced endosomal trafficking of βarr2 using a NanoBiT-based assay. We observed a near-complete loss of endosomal trafficking of βarr2 for the cluster2 mutant in agreement with the confocal imaging data presented earlier (**Figure 7I-J**). Considering that CXCR7 effectively recruits both, βarr1 and 2, we also measured agonist-induced response for these mutants in NanoBiT-based βarr1 recruitment and endosomal trafficking assays. We observed a pattern that is qualitatively similar to βarr2 for both, CXCL12 and VUF11207, although the loss of βarr1 recruitment for Thr^352^ mutant was even more pronounced (**Figure S8A-F**). In these experiments, the surface expression of different mutants was optimized to be comparable to the wild-type receptor (**Figure S7A-J**). Taken together, these data suggest that Thr^352^ is a key residue involved in directing βarr recruitment and trafficking for CXCR7.

**Figure 7.**
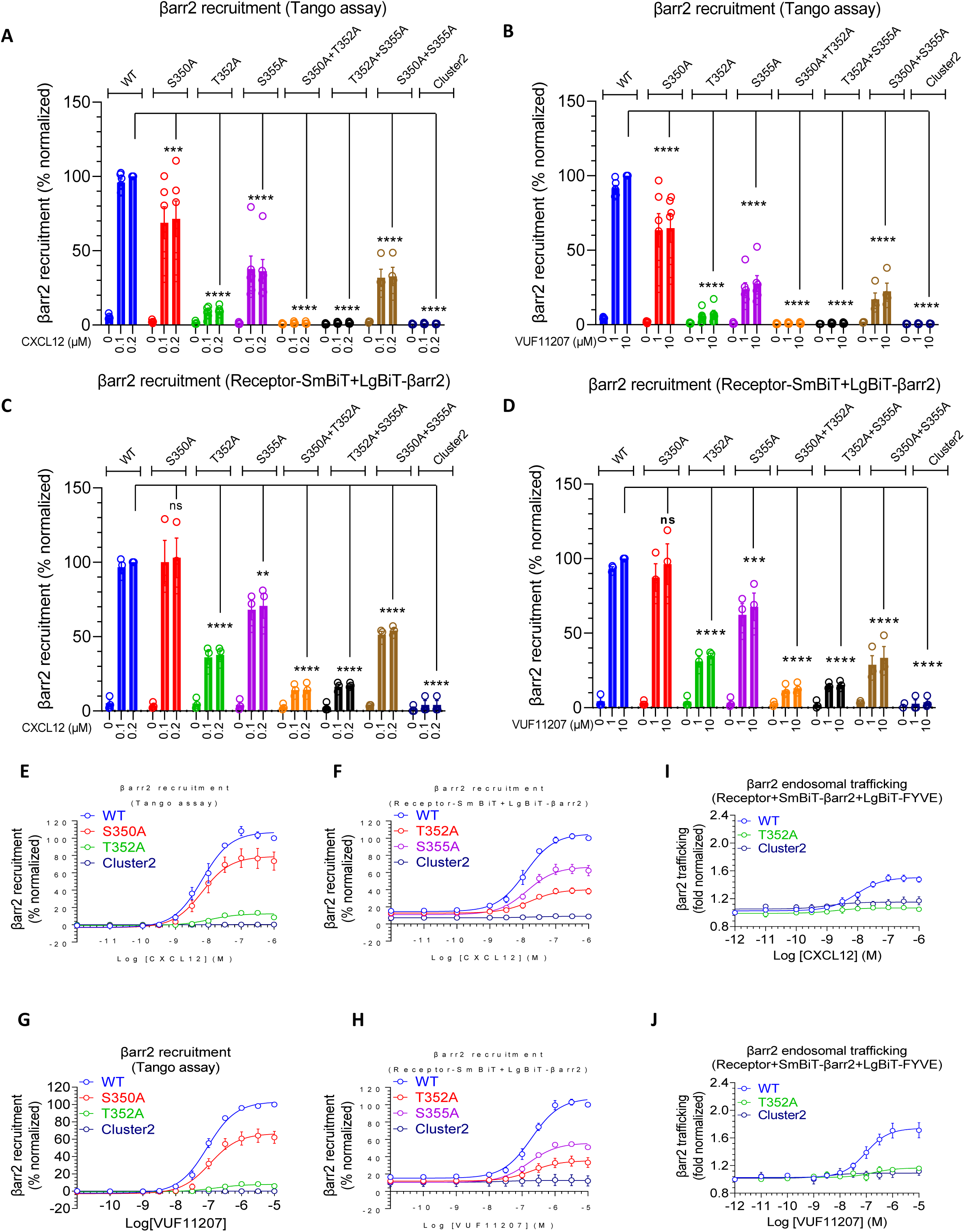
Contribution of the key phosphorylation sites in CXCR7 on βarr recruitment and trafficking. **A-B.** CXCL12- and VUF11207-induced βarr2 recruitment, respectively, to the indicated phosphorylation site mutants of CXCR7 using the Tango assay (mean±SEM; n=6; normalized with luminescence signal at maximal ligand dose for wild-type as 100%, Two-way ANOVA, Tukey’s multiple comparison test, **p<0.01, ***p<0.001, **** p<0.0001, ns= non-significant ). **C-D.** CXCL12- and VUF11207-induced βarr2 recruitment, respectively, to the indicated phosphorylation site mutants of CXCR7 using the NanoBiT assay (mean±SEM; n=3; normalized with luminescence signal at maximal ligand dose for wild-type as 100%, Two-way ANOVA, Tukey’s multiple comparison test, **p<0.01, ***p<0.001, **** p<0.0001, ns= non-significant). **E-H.** Dose response curves of CXCL12- and VUF11207-induced βarr2 recruitment to selected phosphorylation site mutants of CXCR7 in the NanoBiT assay (Receptor-SmBiT+LgBiT-βarr1/2) (mean±SEM; n=6 for panel E,G and n=3 for panel F,H; normalized with luminescence signal for WT at maximal ligand dose as 100%). **I-L.** Dose response curves of CXCL12- and VUF11207-induced βarr2 trafficking to the endosomes for the selected phosphorylation site mutants of CXCR7 in the NanoBiT assay (Receptor+SmBiT-βarr2+LgBiT-FYVE) (mean±SEM; n=3-7 for panel I and J; normalized with the luminescence signal at minimal ligand dose as 1).

## Discussion

The conceptual framework of biased agonism has focused primarily on ligands that induce distinct preferences of transducer-coupling, although biased receptor mutants have also been described for a handful of receptors (Wisler et al., 2014). Chemokine receptors are peculiar in this context as they display a significantly higher degree of ligand promiscuity compared to other prototypical GPCRs, and therefore, may contain several examples of naturally-encoded ligand and receptor bias. Interestingly, a chemokine receptor referred to as the decoy D6 receptor (D6R) has been characterized extensively as an exclusively βarr-coupled receptor (ACR), and it exhibits several important differences compared to its prototypical GPCR counterpart, CCR2 (Pandey et al., 2021). Our data now establish the CXCR4-CXCR7 system as an intriguing example of GPCR-ACR pair, and uncover intrinsic bias encoded at the level of transducer-coupling and functional responses. Moreover, our findings also establish VUF11207 as a highly selective agonist for CXCR7 and demonstrate that its efficacy is comparable to CXCL12, although the potency is slightly weaker. It is also noteworthy that despite a robust recruitment of βarrs, CXCR7 fails to elicit any measurable ERK1/2 phosphorylation in response to either CXCL12 or VUF11207. These data converge with the previous study on D6R, which is also incapable of activating ERK1/2 phosphorylation despite robust interaction with βarrs (Pandey et al., 2021) and further strengthen the emerging notion that βarr-binding does not always translate into ERK1/2 activation at 7TMRs.

The crystal structure of βarr2 in complex with CXCR7pp2 also reveals several intriguing insights and raises tantalizing questions. For example, why does βarr2 undergo a smaller inter-domain rotation compared to V_2_Rpp-bound βarr1 despite a disruption of the three-element interaction and polar core network? While it may reflect a difference between the two isoforms of βarrs, it appears more likely that the number of phosphate groups present on the phosphopeptides may directly influence the inter-domain rotation, possibly through fine-tuning the extent of interactions with βarrs. Going forward, additional structures of βarrs in complex with other phosphopeptides harboring a range of phosphates, derived from different GPCRs and ACRs, may help clarify this. We also acknowledge that the crystal structure contains only the carboxyl-terminus phosphopeptide derived from CXCR7 and lack the transmembrane core of the receptor, which may impart additional changes in βarr conformations. The identification of Thr^352^ as a key site driving CXCR7-βarr recruitment and endosomal trafficking of βarrs presents an interesting parallel to previous studies (Baidya et al., 2020; Chen et al., 2020; Dwivedi-Agnihotri et al., 2020; Zarca et al., 2021). Collectively, these data support the paradigm where the spatial positioning of a single site may be more critical in determining βarr binding and functional outcomes in addition to the phosphorylation bar-code.

In summary, our study provides novel molecular insights into the intrinsic bias encoded in the CXCR4-CXCR7 system, and sheds light on structural mechanism of βarr2 activation and trafficking. These findings represent an important advance to better understand the framework of biased agonism, and intricate details of GPCR-βarr interaction and signaling with potential therapeutic implications.

## Acknowledgements

Research in A.K.S.’s laboratory is supported by the Senior Fellowship of the DBT Wellcome Trust India Alliance (IA/S/20/1/504916) awarded to A.K.S., Science and Engineering Research Board (EMR/2017/003804, SPR/2020/000408, and IPA/2020/000405), Council of Scientific and Industrial Research [37(1730)/19/EMR-II], Indian Council of Medical research (F.NO.52/15/2020/BIO/BMS), Young Scientist Award from Lady Tata Memorial Trust, and IIT Kanpur. A.K.S. is an EMBO Young Investigator and Joy Gill Chair Professor. H.D.-A. is supported by a BioCare grant from DBT (BT/PR31791/BIC/101/1228/2019). The authors thank the staff at Beamline 7A of the Pohang Light Source for their assistance during the X-ray experiments. This work was also supported by Samsung Science & Technology Foundation and Research (SSTF-BA2101-13) and the NRF grants (2015M3D3A1A01064919, 2021R1A2C1004388, 2022R1A2B5B02002529, 2022R1A5A6000760). A.I. was funded by the LEAP JP20gm0010004, and the BINDS JP20am0101095 from the Japan Agency for Medical Research and Development (AMED); Japan Society for the Promotion of Science (JSPS) KAKENHI grants 21H04791, 21H05113, JPJSBP120213501 and JPJSBP120218801; FOREST Program JPMJFR215T and JST Moonshot Research and Development Program JPMJMS2023 from Japan Science and Technology Agency (JST); The Uehara Memorial Foundation; and Daiichi Sankyo Foundation of Life Science. This work was also supported by a grant from the Canadian Institutes of Health Research (CIHR) (MOP-74603) to S.A.L., and Y.C. is supported by a doctoral training scholarship from the Fonds de recherche santé Québec. The research work in XC’s laboratory is supported by a grant from the National Science Foundation of China (21272029). G.D.L. is supported by United States NIH grants R01CA238023, U24CA237683, R01CA238042, U01CA210152, R33CA225549, and R37CA222563. K.E.L. is supported by NIH grant R50CA221807.

## Authors’ contribution

PS performed the cAMP assays, βarr recruitment and trafficking assays, ERK1/2 phosphorylation assay, site-directed mutagenesis with help from DS, SP, HD-A, and confocal imaging with help from MB; H-JY, YY, JJ and KM carried out X-ray crystallography and data collection, CCC and KK performed the G-protein dissociation and βarr recruitment in GRK knock-out cells under the supervision of AI; XR synthesized VUF11207 under the supervision of XC, RB helped in solving the crystal structure and contributed to figure preparation with JM, YC carried out the BRET experiment under the supervision of SAL; PK performed the Calcium assay with the guidance from PNY; MKY, MC and AR carried out Fab30 co-IP and limited proteolysis experiments; KEL generated cell lines used for migration experiments performed by Y-CC under supervision of GDL; HHL supervised and managed the crystallography part of the project; AKS supervised and managed the overall project; all authors contributed to data analysis, interpretation and manuscript writing.

## Accession number

The crystallographic coordinate of the βarr2-CXCR7pp2 complex has been deposited in the RCSB Protein Data Bank with accession number (7XYP).

## Data availability statement

All the relevant data are included in the manuscript. Any additional information can be obtained from the corresponding authors upon reasonable request.

## Materials and methods

### General reagents, plasmids, and cell culture

Most of the general reagents were purchased from Sigma Aldrich unless mentioned otherwise. Dulbecco′s Modified Eagle′s Medium (DMEM), Fetal-Bovine Serum (FBS), Dulbecco’s Phosphate buffer saline (PBS), Trypsin-EDTA, Hank’s balanced salt solution (HBSS), and penicillin-streptomycin solution were purchased from Thermo Fisher Scientific. HEK293 cells (ATCC) were maintained in DMEM (Gibco, Cat. no. 12800-017) supplemented with 10% FBS (Gibco, Cat. no. 10270-106) and 100 U ml^-1^ penicillin (Gibco, Cat. no. 15140122) and 100 μg ml^-1^ streptomycin (Gibco, Cat. no. 15140-122) at 37°C in 5% CO_2_. The cDNA coding region for CXCR4 and CXCR7 were cloned in pcDNA3.1 with an N-terminal FLAG tag. PRESTO Tango assay constructs were acquired from Addgene (Cat. no. 1000000068) while the Tango assay constructs were generated based on previously described framework (Barnea et al., 2008). CXCR7 phosphorylation site mutants were generated by site-directed mutagenesis using Q5 Site-Directed Mutagenesis Kit (NEB, Cat. no. E0554S). For the NanoBiT assay, receptor constructs with carboxyl-terminus SmBiT were generated and other constructs have been described previously^18^. All constructs were verified by DNA sequencing (Macrogen). Recombinant CXCL12 was purchased from PeproTech (Cat. no. 300-28A), and VUF11207 was from Sigma (Cat. no. SML0669). In addition, we also synthesized VUF11207 following a previously published protocol (Wijtmans et al., 2012) and used it in some experiments. The antibodies used in this study were HRP-conjugated anti-FLAG M2 (1:5000; Sigma-Aldrich, Cat. no. A8592), anti-pERK1/2 (1:5000; Cell Signaling Technology, Cat. no. 9101), and anti-tERK1/2 (1:5000; Cell Signaling Technology, Cat. no. 9102). CXCR7pp2 was synthesized at the Tufts University Core Facility.

### Synthesis of VUF11207

The synthesis of VUF11207 is described below in the schematic. Aldol condensation of 2-fluorobenzaldehyde with propionaldehyde under basic condition gave (E)-3-(2-Fluorophenyl)-2-methylacrylaldehyde (1) in 72% yield. In the presence of acetic acid, aldehyde 1 reacted with 2- (1-methylpyrrolidin-2-yl) ethanamine in methanol, and the resulting imine was reduced with NaBH(OAc)3 to generate (E)-3-(2-fluorophenyl)-2-methyl-N-(2-(1-methylpyrrolidin-2-yl)ethyl) prop-2-en-1-amine (2) in 50% yield. Under the standard amide coupling conditions (EDCI/HOBt/DIEA), amine 2 was coupled with 3,4,5-trimethoxybenzoic acid to afford VUF11207 with 67% yield. The structures of target compound and the intermediates were confirmed by their spectral properties. A full description of VUF11207 synthesis and characterization will be published separately.

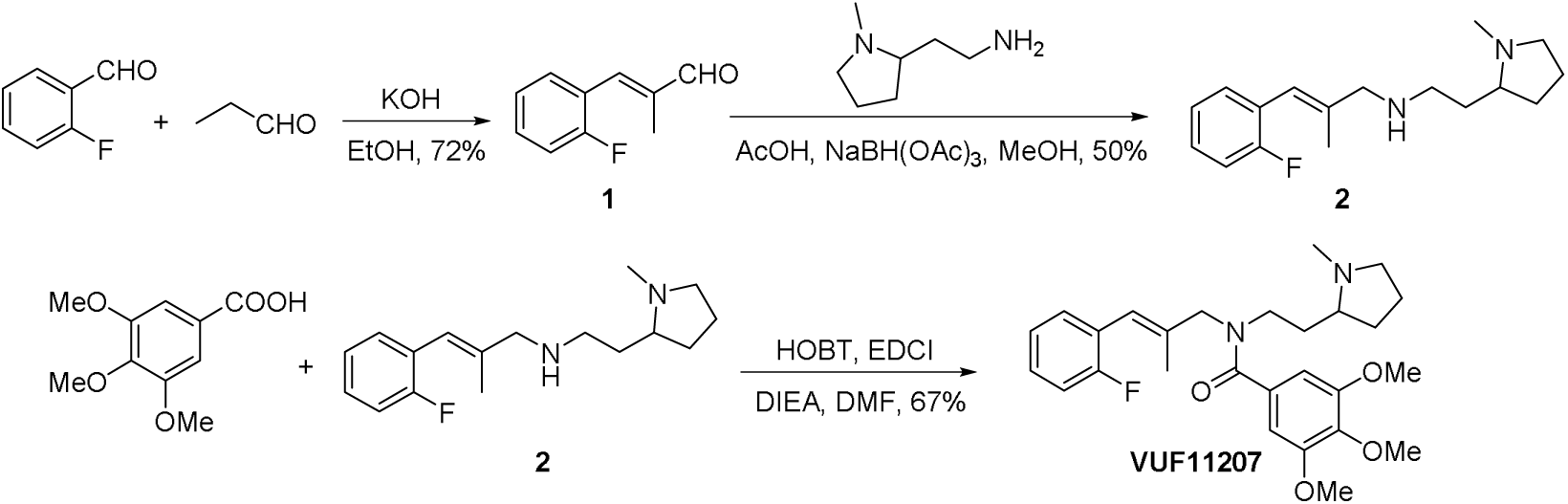

### NanoBiT-based G-protein dissociation assay

Ligand-induced G-protein activation was measured using a previously described NanoBiT-based G-protein dissociation assay(Inoue et al., 2019). Briefly, a NanoBiT-G-protein consisting of LgBiT-tagged Gα subunit and SmBiT-tagged Gγ2 subunit along with the untagged Gβ1 subunit were co-expressed with the indicated receptor constructs and ligand-induced change in luminescence signal was measured. Typically, HEK293A cells (Thermo Fisher Scientific) were transfected with a plasmid mixture consisting of 100 ng LgBiT-Gα, 500 ng Gβ1, 500 ng SmBiT-Gγ2 (C68S) with either 50 ng of CXCR4 plasmid or 3.5 μg of CXCR7. The receptor constructs used in this assay contain N-terminal HA signal sequence, FLAG tag and a flexible linker sequence. To enhance NanoBiT-G-protein expression for Gs, Gq and G12/13, 100 ng of RIC8B plasmid (isoform 2; for Gs) or RIC8A (isoform 2; for Gq, G12, and G13) were also co-transfected. 24 h post-transfection, cells were harvested with EDTA-containing PBS, centrifuged, and suspended in 2 ml of HBSS (Gibco, Cat. no. 14065-056) containing 0.01% bovine serum albumin (BSA fatty acid–free grade, SERVA) and 5 mM 4-(2-hydroxyethyl)-1-piperazineethanesulfonic acid (HEPES), pH 7.4 (assay buffer). Afterwards, cells were dispensed in a white 96-well plate (80 μl well^-1^), incubated with 20 μl of 50 μM coelenterazine (Carbosynth, Cat. no. EC175526), and 2 h later, baseline luminescence was measured (SpectraMax L, Molecular Devices). Subsequently, 20 μl of 6X agonist, serially diluted in the assay buffer, were manually added and the plate was immediately read for the second measurement in a kinetic mode. Luminescence counts recorded from 3-5 min post-agonist addition were averaged, corrected with the baseline signals, normalized with respect to vehicle control plotted using the GraphPad Prism 9 software.

### GloSensor-based cAMP assay

In order to assess agonist-induced coupling of Gαs and Gαi, we used GloSensor-based cAMP assay following a protocol described previously (Kumar et al., 2017). In brief, HEK293 cells were co-transfected with FLAG-tagged receptor constructs and luciferase-based 22F cAMP biosensor (Promega, Cat. no. E2301) using polyethylenimine (PEI) linear (Polysciences, Cat. no. 19850) at a ratio of 1:3 (DNA:PEI linear) as transfection reagent. After 14-16 h of transfection, cells were detached from the plates, resuspended in assay buffer (1XHBSS, 20 mM HEPES, pH7.4) containing D-luciferin (0.5 mg ml^-1^, GoldBio, Cat. no. LUCNA-1G) and seeded into 96 well white plates (Corning) at a density of 2×10^5^ cells well^-1^ in a volume of 100 μl. After an incubation of 1.5 h at 37°C and 30 min at room-temperature, baseline luminescence readings were recorded. For Gαs assay (**Figure 1E**, middle panel), ligands prepared in the assay buffer were added at indicated final concentration after baseline readings while for Gαi assay (**Figure 1E**, left panel and **Figure 2L**), 5 μM forskolin (Sigma, Cat. no. F6886) was added to the cells and luminescence readings were recorded till they stabilized (5-10 cycles) followed by ligand addition. The change in luminescence signal was recorded using a microplate reader (Victor X4; Perkin Elmer) for 30 min, and data were normalized as indicated in the respective figure legends and plotted using GraphPad Prism 9 software. For the experiments presented in **Figure 1E** (left panel), 0.25 μg of CXCR4 and 5 μg of CXCR7 were used along with 2 μg of 22F plasmid. For the Gαs-coupling assay (**Figure 1E**, middle panel), 0.5 μg of CXCR4, 0.5 μg of V_2_R and 4 μg of CXCR7 plasmids were used along with 3 μg of 22F plasmid. For the experiments presented in **Figure 2L**, 3.5 μg of CXCR1, CXCR3, and CXCR4 plasmids were used while 5 μg of plasmids were used for CXCR2, CXCR5, CXCR6, and CXCR7, along with 2 μg of 22F plasmid.

### Calcium flux assay

HEK293T cells were transiently transfected with pGP-CMV-GcAMP6s (Ca^2+^ Sensor plasmid, Addgene, Cat. no. 40753; 4 µg), 5HT_2c_ receptor (as positive control, cDNA.org, Cat. no. HTR02CTN00; 4 µg) or CXCR4/CXCR7 receptor (4 µg) using PEI max in a ratio of 1:4 (DNA:PEI max) and plated at a density of 50,000 cells well^-1^ in black optical bottom plate in complete DMEM media (10% FBS). After 14-16 h of transfection, media from the plate was aspirated and 100 μl of Ca^2+^/Mg^2+^ free HBSS buffer (pH 7.2) was added, cells were further incubated at 37°C for 10 min in the Flex Station 3 (Molecular Devices) before the assay was initiated. Ligand induced change in relative fluorescence unit (RFU) was measured at excitation 485 nm and emission 525 nm (cut off 515 nm) with the settings of 6 reads well^-1^. Basal fluorescence of each well was recorded for 15 s, and then 20 µl of 6X concentration of each agonist (Serotonin and VUF11207) as indicated was added using robotic pipetting of FlexStation system and RFU was recorded at 2 s interval for a total of 135 s. The changes in RFU (ΔRFU) for each treatment group was calculated by subtracting the average basal response (RFU before ligand addition) from RFU of each well at each time points after ligand addition. ΔRFU for each ligand was plotted and analyzed using GraphPad Prism 9.

### Receptor surface expression

In order to measure the surface expression of the receptors in various assays, we used a previously described whole cell-based surface ELISA assay(Pandey et al., 2019). Transfected cells from the corresponding assays were seeded into a 24-well plate pre-coated with 0.01% poly-D-Lysine at a density of 2×10^5^ cells well^-1^ and incubated at 37°C for 24 h. Afterwards, cells were washed once with ice-cold 1XTBS, fixed with 4% PFA (w/v in 1XTBS) on ice for 20 min, washed again three times with 1XTBS, and blocked at room temperature for 1.5 h with 1% BSA prepared in 1XTBS. Subsequently, the cells were incubated with anti-FLAG M2-HRP antibody (Sigma, Cat. no. A8592) (1:2000, 1.5 h at room temperature) followed by three washes in 1% BSA and incubation with TMB-ELISA substrate (Thermo Fisher Scientific, Cat. no. 34028) until the light blue color appeared. The signal was quenched by transferring 100 μl of the colored solution to another 96-well plate containing 100 μl of 1 M H_2_SO_4_, and the absorbance was measured at 450 nm. For normalization, TMB substrate was removed, cells were washed twice with 1XTBS, and incubated with 0.2% (w/v) Janus Green (Sigma, Cat. no. 201677) for 15 min at room temperature. The excess stain was removed by washing the cells with water followed by addition of 800 μl of 0.5 N HCl in each well, and 200 μl of this solution was transferred to a 96-well plate for measuring the absorbance at 595 nm. The signal intensity was normalized by calculating the ratio of A450/A595 values and plotted using the GraphPad Prism 9.

For the NanoBiT-based G-protein dissociation assay and βarr recruitment in GRK knock-out cells, surface expression of the receptors was measured using flow-cytometry based assay following a previously described protocol (Pandey et al., 2021). Briefly, a small amount of HEK293A cells from the corresponding assays were harvested with 0.5 mM EDTA-containing PBS and transferred to a 96-well V-bottom plate. Cells were fluorescently labeled using anti-FLAG monoclonal antibody (Clone 1E6, FujiFilm Wako Pure Chemicals; 10 μg ml^-1^ diluted in 2% goat serum+2 mM EDTA-containing PBS) followed by incubation with Alexa Fluor 488-conjugated goat anti-mouse IgG secondary antibody (Thermo Fisher Scientific; 10 μg ml^-1^). Subsequently, the cells were washed with PBS, resuspended in 2 mM EDTA-containing PBS, filtered through a 40 μm filter and the fluorescent intensity of single cells was quantified using a flow cytometer. Fluorescent signal from Alexa Fluor 488 was recorded and analyzed using the FlowJo software. Mean fluorescence intensity from about 20,000 cells per sample were used for analysis.

### Tango assay for βarr recruitment

In order to assess the βarr2 recruitment to indicated receptors, Tango assay was used following the previously published protocol(Dogra et al., 2016). HTLA cells were transfected with indicated receptor constructs and 24 h post-transfection, cells were trypsinized, resuspended in complete DMEM, and seeded into 96-well white plates at a density of 1X10^5^ cells well^-1^. After another 24 h, cells were stimulated with the indicated dose of ligands and incubated at 37°C for additional 7-8 h. Afterwards, the culture media was changed with assay buffer (1XHBSS, 20 mM HEPES, pH 7.4 and 0.5 mg ml^-1^ D-luciferin). Luminescence readings were measured in a microplate reader (Victor X4; Perkin Elmer), data were corrected for baseline luminescence readings, normalized as mentioned in the corresponding figure legends and plotted using GraphPad Prism 9. For the data presented in **Figure 2C, 2H** and **S2H**, PRESTO Tango constructs were used as described previously (Kroeze et al., 2015). For the data presented in **Figure 2B, 2D** and **2K**, Tango assay constructs were generated by engineering a TEV protease cleavage site and tTa transcription factor at the end of the receptor coding sequence in pcDNA3.1 vector backbone.

### Microfluidic chemotaxis assay

We quantified chemotaxis using a previously described microfluidic device that tracks movement of single cells toward a gradient (Chen et al., 2015) and we used MDA-MB-231 human breast cancer cells (purchased from the ATCC, Manassus, VA, USA) stably transduced with CXCR7 fused to GFP (Luker et al., 2010). Briefly, we introduced MDA-MB-231 cells stably transduced with CXCR7 fused to GFP into the device at a concentration of 1 x 10^5^ cells/ml in complete DMEM medium with 10% serum and 1% GlutaMAX. After allowing cells to adhere for 10 minutes, we replaced medium in the seeding port with serum-free DMEM. We added a chemoattractant, 100 ng ml^-1^ CXCL12-α (R&D Systems) and/or 100 nM VUF11207 (Cayman Chemical) in serum-free DMEM with 0.1% Probumin (Millipore), to the opposite side of the device. We quantified chemotaxis of single cells after 16 h in the device.

### NanoBiT assay for βarr recruitment

Agonist-induced βarr1/2 recruitment for CXCR4 and CXCR7 was also measured using NanoBiT-based assay following the protocol described earlier (Pandey et al., 2021). Briefly, HEK293 cells were transfected with CXCR4 (1 µg) and CXCR7 (7 µg) harboring carboxyl-terminus fusion of SmBiT and βarr1/2 constructs (2 µg) with N-terminal fusion of LgBiT. The cells were stimulated with varying doses of indicated ligands followed by measurement of luminescence signal using a multimode plate reader for 10-15 cycles and average data from 5^th^ to 10^th^ cycle are used for analysis and presentation. In order to evaluate the contribution of different GRKs in βarr recruitment to CXCR7, we used previously described GRK knock-out cell lines (Kawakami et al., 2022) and the NanoBiT assay as described above.

### ERK1/2 MAP kinase phosphorylation assay

Agonist-induced ERK1/2 MAP kinase phosphorylation was measured following the previously published protocol (Kumari et al., 2019). Briefly, HEK293 cells were transfected with CXCR4 (0.25 µg), CXCR7 (4 µg) or empty vector (pcDNA3.1; 7 µg), and 24 h post-transfection, they were seeded into a 6-well plate at a density of 1×10^6^ cells well^-1^. Subsequently, the cells were serum starved for 12 h followed by agonist-stimulation as indicated in the corresponding figure legends. Afterwards, the cells were harvested, lysed in 2XSDS loading buffer, heated at 95°C for 15 min followed by centrifugation at 15000 rpm for 15 min. 10 μl of lysate was then separated by SDS-PAGE and ERK1/2 bands were detected by Western blotting using corresponding antibodies (rabbit phospho-ERK1/2 antibody, 1:5000 dilution; rabbit total ERK1/2 antibody, 1:5000 dilution; anti-rabbit HRP-coupled secondary antibody, Genscript, Cat. No. A00098, 1:10000 dilution). ECL solution from Promega (Cat. no. W1015) was used as a substrate for the HRP, and the signals were developed using ChemiDoc (BioRad). The signals were quantified using densitometry in BioRad Image Lab software, normalized as indicated in the figure legend, and data were plotted using GraphPad Prism 9. For the experiments presented in **Figure 3C-D**, cells were pre-treated with 10 μM of AMD3100 for 30 min prior to CXCL12 stimulation, and for the experiments presented in **Figure S3C-D**, cells were pre-treated with Pertussis toxin (100 ng μl^-1^) for 12 h during serum starvation step.

### BRET assay for βarr2 conformational change

Intramolecular FlAsH-based BRET sensors were used to monitor the conformational changes in βarr2 following the previously published protocol (Lee et al., 2016; Nuber et al., 2016). In brief, HEK293SL cells were seeded at a density of 1.5×10^5^ cells well^-1^ in 6-well plates and transfected with the indicated receptor constructs along with the βarr2-FlAsH sensors using calcium phosphate. 24 h post-transfection, cells were detached and seeded into poly-ornithine-coated 96-well white plates at a density of 2.5×10^4^ cells well^-1^. After another 24 h, cells were washed and incubated with Tyrode’s buffer (140 mM NaCl, 2.7 mM KCl, 1 mM CaCl_2_, 12 mM NaHCO_3_, 5.6 mM D-glucose, 0.5 mM MgCl_2_, 0.37 mM NaH_2_PO_4_, 25 mM HEPES, pH 7.4) for 1 h at room temperature. Subsequently, FlAsH reagent solution was prepared by mixing 1.75 μl of FlAsH-EDT2 stock reagent with 3.5 μl of 25 mM EDT solution in DMSO and left for 10 min at room temperature. 100 μl of Tyrode’s buffer was added to this mixture followed by an additional incubation for 5 min at room temperature and then the volume was adjusted to 5 ml with Tyrode’s buffer. Cells were incubated with 60 μl of the labeling solution for 1 h at 37°C followed by washing with BAL wash buffer and Tyrode’s buffer. Finally, 90 μl of Tyrode’s buffer was added to each well and the plate was incubated at 37°C for 1 h before ligand stimulation. Coelenterazine H was added at a final concentration of 2 μM, cells were stimulated with 100 nM CXCL12 and six consecutive BRET measurements were taken using a Victor X; PerkinElmer plate reader with a filter set (center wavelength/band width) of 460/25 nm (donor) and 535/25nm (acceptor). BRET ratios (intensity of light emitted by the acceptor/intensity of light emitted by the donor) were calculated and net-BRET ratio was determined after subtracting the background BRET ratio i.e. the difference between the FlAsH-EDT2-labeled BRET ratio and the unlabeled condition. The difference of the net-BRET ratio for ligand-stimulated condition vs. vehicle-treatment was plotted using GraphPad Prism 9.

### Confocal microscopy

In order to visualize βarr2 trafficking upon stimulation of wild-type and phosphorylation site mutants, we used confocal microscopy following the previously published protocol (Dwivedi-Agnihotri et al., 2020). HEK293 cells were co-transfected with 5 µg of indicated receptor constructs and 2 µg of βarr2-mYFP plasmid using PEI. 24 h post-transfection, cells were trypsinized, seeded onto poly D-lysine coated glass-bottom confocal dishes (SPL, Cat. No. 100350) at a density of 1×10^6^ per dish. After overnight incubation, cells were starved in FBS deficient DMEM for 4 h and visualized under the microscope. Cells were stimulated with either CXCL12 (100 nM) or VUF11207 (1 μM) and live-cell imaging was performed on Zeiss LSM 710 NLO confocal microscope. A diode laser at a 488 nm laser line was used for exciting mYFP, and the emitted signal was detected with a 32xarray GaAsP descanned detector (Zeiss). Images were further processed with Zen software suite (Zeiss).

### βarr2 expression and purification

CXCR7 phosphopeptide (CXCR7pp2) for crystallization and ITC experiments were obtained from NovoPep. The *Rattus norvegicus* wild-type βarr2 (1–356) were inserted into expression vector pET-28a. The plasmids were transformed into *Escherichia coli* BL21(DE3)pLysS cells (Invitrogen), and cells harboring the plasmids were grown at 37°C until the OD_600_ reached 0.7-1.0 in Luria Bertani (LB) broth containing 70 μg ml^-1^ chloramphenicol and 30 μg ml^-1^ kanamycin. Further, 0.1 mM isopropyl-β-D-1-thiogalactopyranoside (IPTG) was used to induce protein expression in the cells, after which the cells were incubated for 16 h at 16°C. For isolation of βarr2 protein fused to an N-terminal His6-tag, cells were harvested by centrifugation at 5000 rpm at 4°C for 10 min, and the pellet was resuspended in ice-cold buffer A (20 mM Tris-HCl, pH 8.0, 500 mM NaCl, and 5 mM Imidazole) containing 1 mM phenylmethanesulfonylfluoride (PMSF). The cells were lysed using a microfluidizer (Microfluidics, Westwood, MA, USA), and the lysed cells were centrifuged at 15,000 rpm (Vision V506CA rotor) at 4°C for 30 min to separate the supernatant and cell debris. The supernatant was applied to a Ni-sepharose affinity column (GE Healthcare, Little Chalfont, UK) pre-equilibrated with buffer A. Initially, the column was washed extensively with buffer A, after which the protein was eluted using buffer A containing a gradient of imidazole concentrations from 100 mM to 1 M. The eluates were desalted into buffer B (20 mM Tris-HCl, pH 8.0 and 5 mM β-mercaptoethanol) containing 100 mM NaCl by using desalting column (GE Healthcare) and further purified by affinity chromatography with HiTrap heparin column (GE Healthcare). The proteins were eluted using buffer B containing 1 M NaCl in heparin column. Further purification was performed by gel filtration on a HiLoad 16/60 Superdex 200 prep-grade column (GE Healthcare), which was equilibrated with buffer B containing 200 mM NaCl. The homogeneity of the all purified protein was assessed by polyacrylamide gel electrophoresis in the presence of 0.1% (w/v) sodium dodecyl sulfate. For crystallization, protein solutions were concentrated to about 12 mg m^l-1^ using a centricon centrifugal filter unit (Sartorius Stedim). The protein concentration was estimated by measuring the absorbance at 280 nm.

### Isothermal Titration Calorimetry (ITC)

ITC experiments were performed using Affinity ITC instruments (TA Instruments, New Castle, DE, USA) at 298K. 200 µM of βarr2 (1–356), which was prepared in a buffer containing 20 mM HEPES pH 7.0 and 200 mM NaCl was degassed at 298 K prior to measurements. Using amicro-syringe, 2 µl of 1 mM CXC7pp2 solutions were added at intervals of 200 s to the βarr2 solution in the sample cell with gentle stirring.

### Crystallization and data collection

Before crystallization, βarr2 (1–356) (12 mg ml^-1^) in buffer B containing 200 mM NaCl and CXCR7pp2 peptide (70 mg ml^-1^) in 150 mM Tris (pH 8.0) were mixed in a 7:1 volume ratio and incubated at 4°C for 1 h. Crystals were grown at 22°C using sitting-drop vapor diffusion by mixing 1 μl of the protein complex solution with 1 μl of 20% (w/v) PEG3350, 0.2 M ammonium acetate, and 0.1 M bis-Tris (pH 5.5). Crystals were cryoprotected by soaking in N-Paratone oil (Sigma-Aldrich) and flash frozen in liquid nitrogen. X-ray diffraction data were collected at 100K in 1° oscillations at the beamline BL-7A of PLS (Supplementary Table 1). Raw X-ray diffraction data were processed and scaled using the program suit HKL200022. Data collection statistics are summarized in (Supplementary Table 1).

### Co-immunoprecipitation (co-IP) assay

Co-IP was performed to evaluate the interaction between V_2_Rpp or CXCR7pp2 with βarr2 in the presence of Fab30. 5 μg of purified βarr2 was activated with a 10-fold molar excess of corresponding phospho-peptides for 1 h at room temperature (25°C) in binding buffer (20 mM HEPES, pH7.4, 100 mM NaCl). Thereafter, the activated βarr1 was incubated with 2.5 μg of purified Fab30. Subsequently, 20 μl of pre-equilibrated Protein L beads (GE Lifesciences, Cat. no. 17547802) were added to the reaction mixture and incubated for an additional 1 h at room temperature, which was followed by extensive washing (3–5 times) with binding buffer + 0.01% LMNG. Elution was taken with 2X SDS loading buffer. Interaction of Fab30 with βarr2 in presence of phospho-peptides was visualized using Coomassie staining of the gels. Band intensity was analyzed by ImageJ gel analysis software.

### Structure determination and refinement

Intensity statistics of the data collected against the βarr21–356-C7pp2 complex crystals from Phenix Xtriage suggested possible twinning, related by pseudo-merohedral twin operators -h, - k, l. The crystal structure of the βarr2 (1–356)-CXCR7pp2 complex was determined by molecular replacement using the monomeric coordinates of rat βarr2 (1–356)-CXCR7pp1 (PDB ID: 6K3F) as the search model. Molecular replacement with Phaser (McCoy et al., 2007) within the CCP4 suite (Potterton et al., 2003; Potterton et al., 2018) yielded a solution containing 6 molecules in the asymmetric unit, with a Vm of 2.21 (44.33% solvent content). Density modification was performed with Resolve within the Phenix suite (Adams et al., 2010), followed by model building with COOT (Emsley and Cowtan, 2004) and refinement with Phenix_refine (Potterton et al., 2018). Although the best diffraction spots reached 1.99 Å, we noticed that the model generated based on molecular replacement did not refine well despite considering twin law (-h, -k, l) during refinement in Phenix. Although the Rfactor and Rfree of the model converged to 23% and 27% respectively, there was overall poor geometry in terms of Ramachandran and rotamer outliers and high clash score. This model could not be improved further even after several rounds of manual rebuilding in the clear electron density and successive refinement with the twin law. Therefore, in order to get a better model in terms of geometry and minimize outliers, we truncated the data to 2.5Å and 2.8Å, respectively, and re-refined the model against the corresponding truncated datasets. Model building and refinement with the 2.8Å truncated dataset yielded a better model with reasonable geometry and minimized Ramachandran outliers although the Rfactor and Rfree were on the higher side (28% and 33.2%, respectively). Data collection and refinement statistics have been included as Supplementary Table 1. The final model contains chains A, B, C, D, E, and F corresponding to βarr2 (1–356), and chains U, V, W, X, Y, and Z corresponding to CXCR7pp2 molecules.

### βarr recruitment for the phosphorylation site mutants of CXCR7

The phosphorylation site mutants of CXCR7 as indicated in **Figure 7** and **S7** were generated using site-directed mutagenesis kit followed by βarr recruitment in Tango and NanoBiT assays as described above. Surface expression of the indicated mutants was first optimized to be at comparable levels followed by the Tango and NanoBiT assays. For the Tango assay, HTLA cells were transfected with 7 µg or the wild-type and mutant receptor constructs except the CXCR7^T352A+S355A^ for which, 5 μg DNA was transfected. Cells were treated with indicated concentration of agonists for 8 h at 37°C followed by the addition of luciferin and luminescence measurement. Data normalization is described in the corresponding figure legends.

### NanoBiT assay for βarr trafficking

Agonist-induced βarr1/2 trafficking to the endosomes was monitored using NanoBiT assay following the same protocol as described above for βarr recruitment except that the receptor constructs were not tagged with SmBiT. Instead, N-terminal SmBiT-tagged βarr1/2 constructs and N-terminal LgBiT-tagged FYVE constructs were used for enzyme complementation. The amount of DNA for receptor, βarr1/2 and FYVE was kept as 3 µg, 2 µg and 2 µg, respectively.

