## Supplemental Material for "Molecular insights into intrinsic transducer-coupling bias in the CXCR4-CXCR7 system"

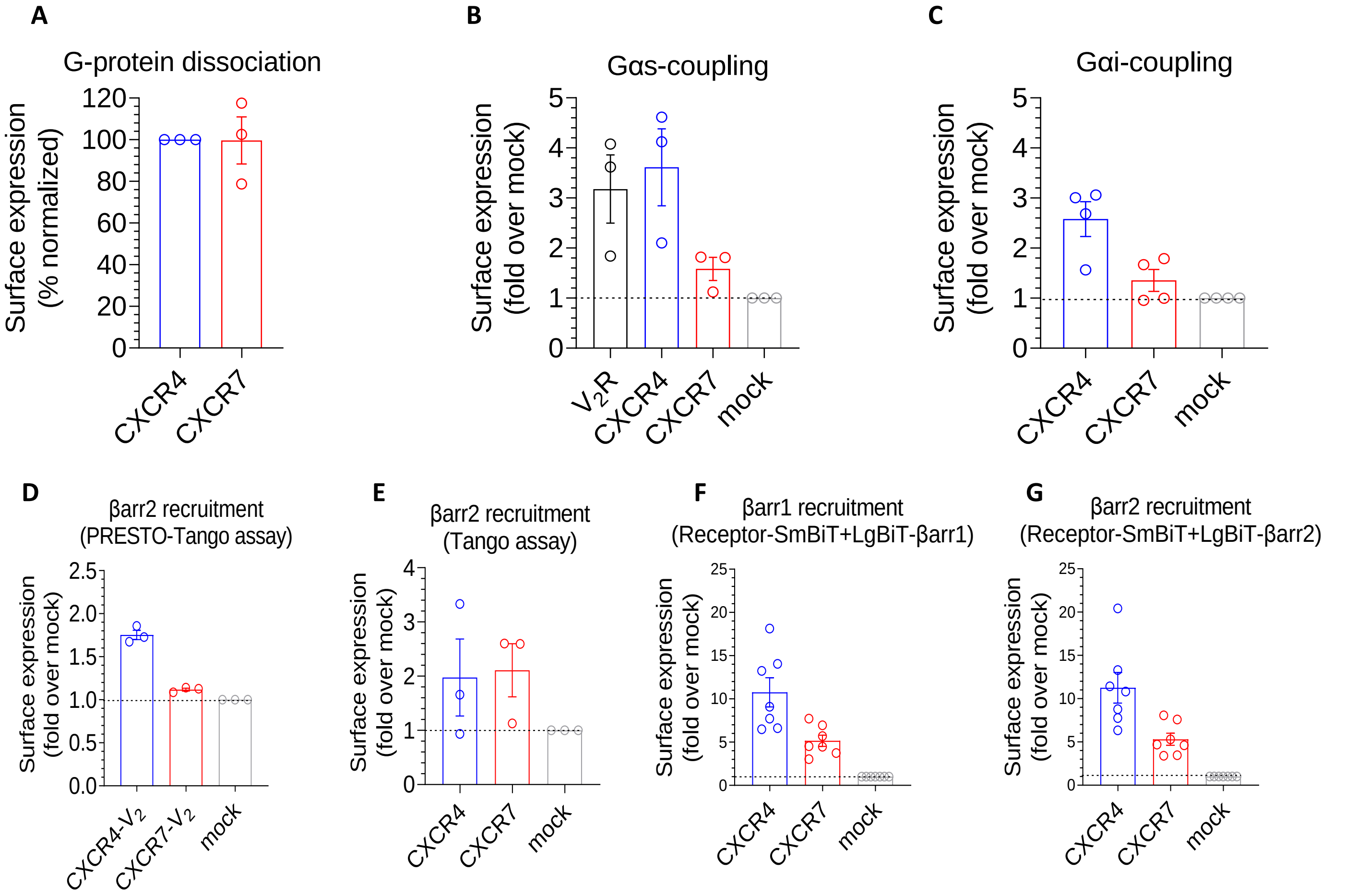


**Figure S1. Surface expression of CXCR4 and CXCR7 in different assays A.** Surface expression of CXCR4 and CXCR7 in the G-protein dissociation assay measured using flow cytometry assay (mean±SEM; n=3 normalized with CXCR4 as 100%). **B-C.** Surface expression of the indicated receptors in cAMP assay as measured using whole cell ELISA (mean±SEM; n=3 for panel B and n=4 for panel C; normalized as fold over mock-transfection). **D-G.** Surface expression of indicated receptors in βarr recruitment assays measured using whole cell ELISA (mean±SEM; n=3 for panel D-E and n=7 for panel F-G; normalized as fold over mock-transfection).

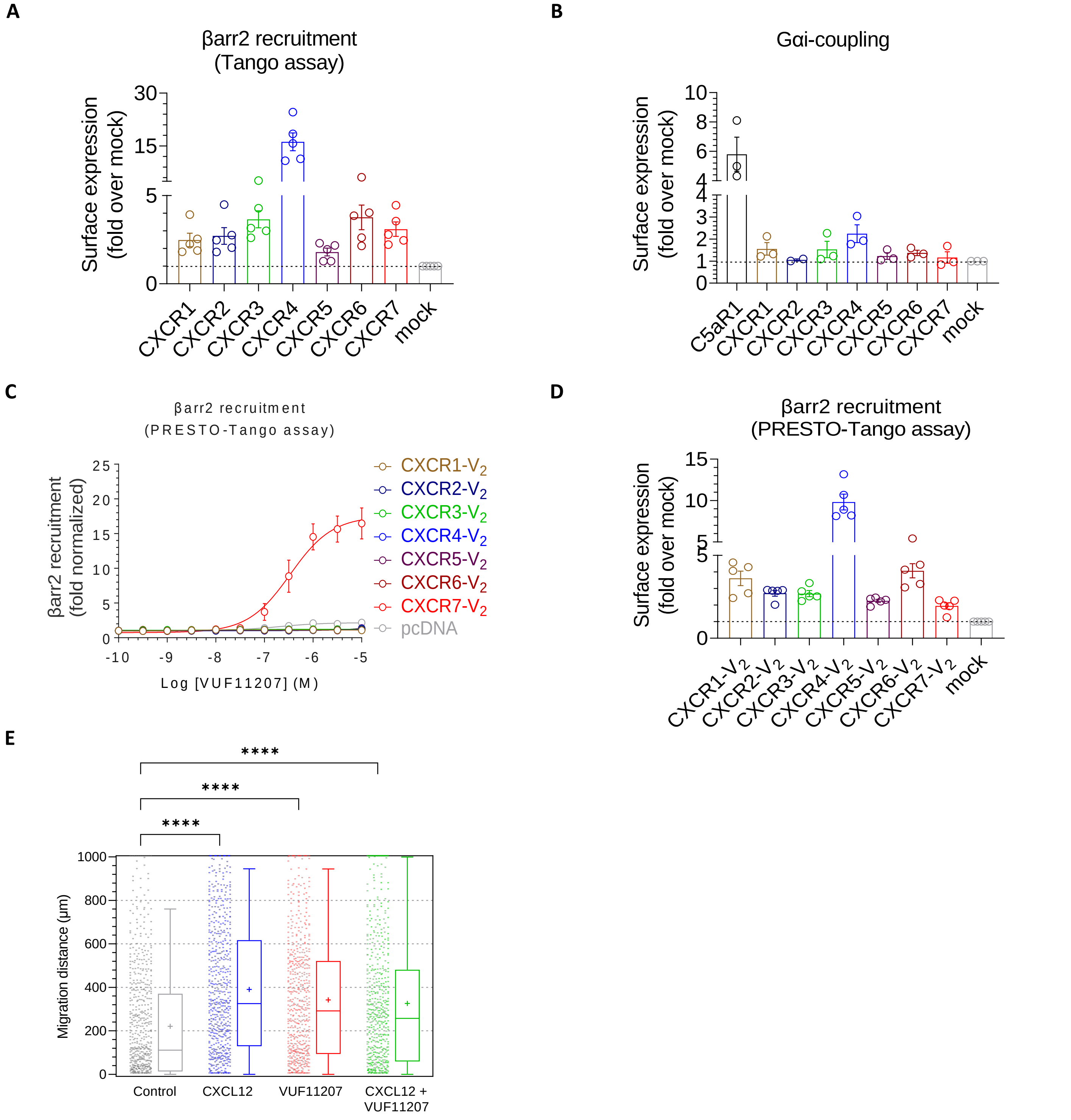


**Figure S2. Surface expression, profiling of VUF11207, and migration of MDA-MB-231 cells. A-B.** Surface expression of indicated receptors were measured in different assays using whole cell ELISA (mean±SEM; n=5 for panel A, n=2-3 for panel B; normalized as fold over mock-transfection). **C.** VUF11207-induced βarr2 recruitment for all the CXC chemokine receptors (CXCR1-7) in the PRESTO Tango assay (mean±SEM; n=5; normalized as fold increase with luminescence signal at the lowest ligand dose). **D.** Surface expression of indicated receptors in the PRESTO Tango assay measured using whole cell ELISA (mean±SEM; n=5 for panel D; normalized as fold over mock-transfection, One-way ANOVA, Sidak's multiple comparison test, **** $p<0.0001$). **E.** Migration of MDA-MB-231 stably expressing CXCR7 in response to indicated ligands. The experiment was carried out using a 2D microfluidic device and each point on the graph represents an individual cell.

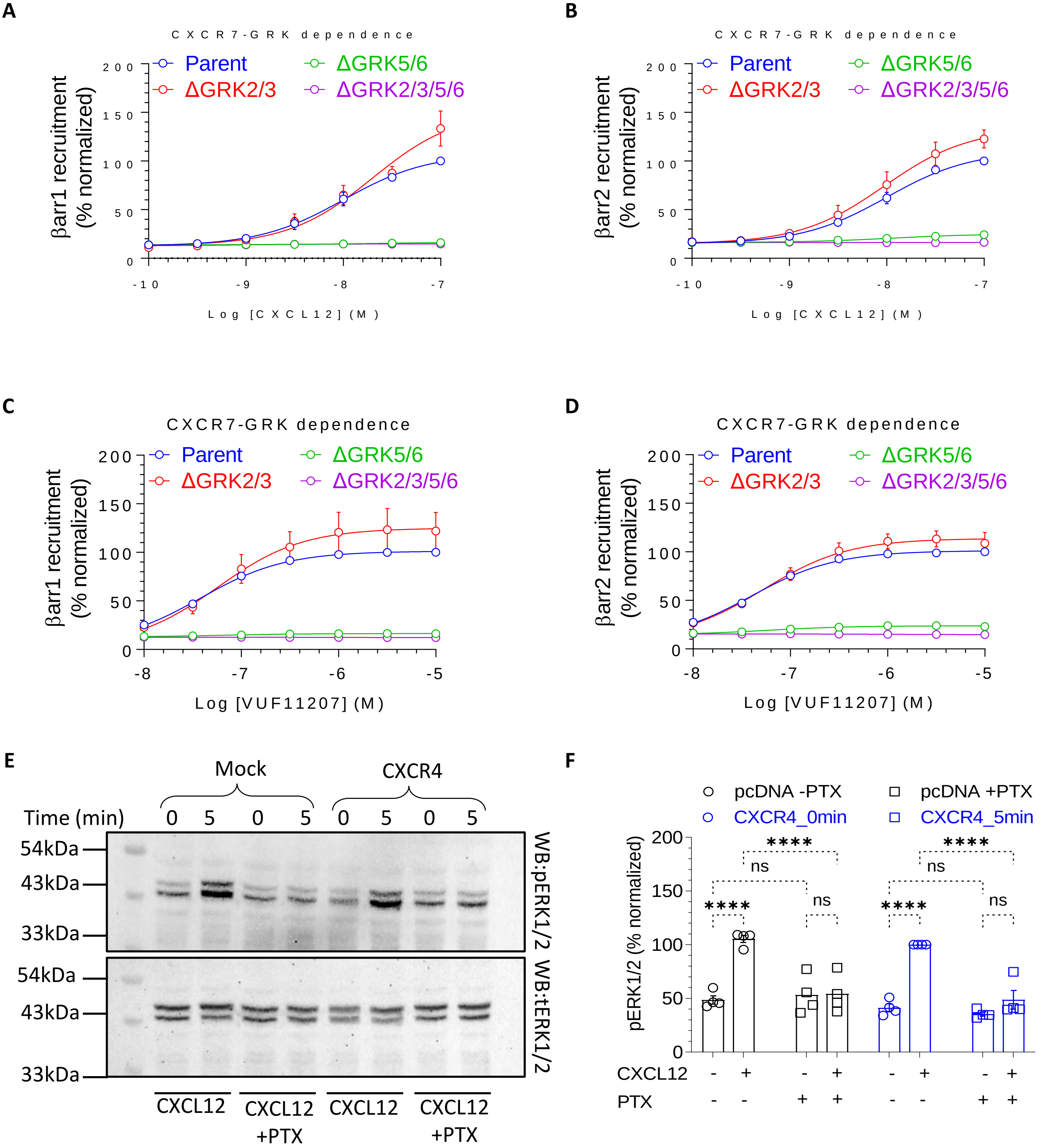


**Figure S3. Effect of PTX treatment on βarr recruitment and ERK1/2 phosphorylation. A-D.** CXCL12- and VUF11207-induced βarr1 and 2 recruitment to CXCR7 in GRK knock-out cells using the NanoBiT assay with co-expression of the catalytic subunit of pertussis toxin (PTX) (Receptor-SmBiT+LgBiT-βarr1/2), respectively (mean±SEM; n=4-5; normalized with luminescence signal at maximal ligand dose as 100%). **E.** CXC12-induced ERK1/2 phosphorylation in HEK239 cells transfected with either empty vector (mock) or CXCR4 plasmid with and without pre-treatment of PTX. **F**. Densitometry-based quantification of data presented in panel E (mean±SEM; n=4; normalized with 5 min stimulation of CXCR4 condition without PTX, Two-way ANOVA, Tukey's multiple comparison test, **** $p<0.0001$, ns= non-significant ).

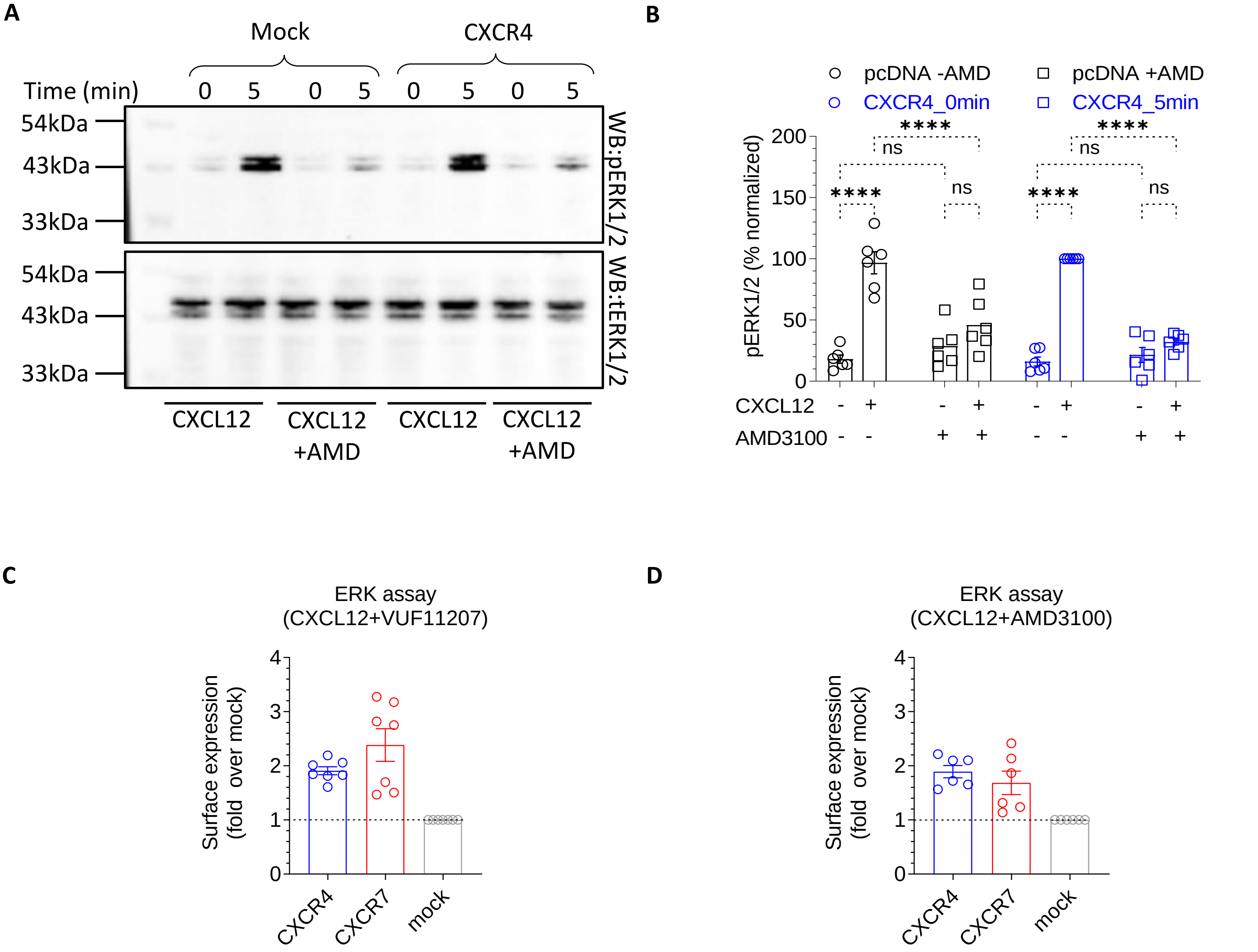


**Figure S4. Pre-treatment with AMD3100 blocks CXCL12-induced ERK1/2 phosphorylation.**
**A.** HEK293 cells transfected with CXCR4 construct or empty vector (mock) were stimulated with CXCL12 with or without AMD3100 pre-treatment followed by detection of ERK1/2 phosphorylation using Western blotting. **B.** Densitometry-based quantification of ERK1/2 phosphorylation (mean±SEM; n=6, normalized with the 5 min signal for CXCR4 as 100%). **C-D.** Surface expression of indicated receptors were measured in different assays using whole cell ELISA (mean±SEM; n=7 for panel C and n=6 for panel D ; normalized as fold over mock-transfection, Two-way ANOVA, Tukey's multiple comparison test, **** p<0.0001, ns= non-significant ).

**A** **B**

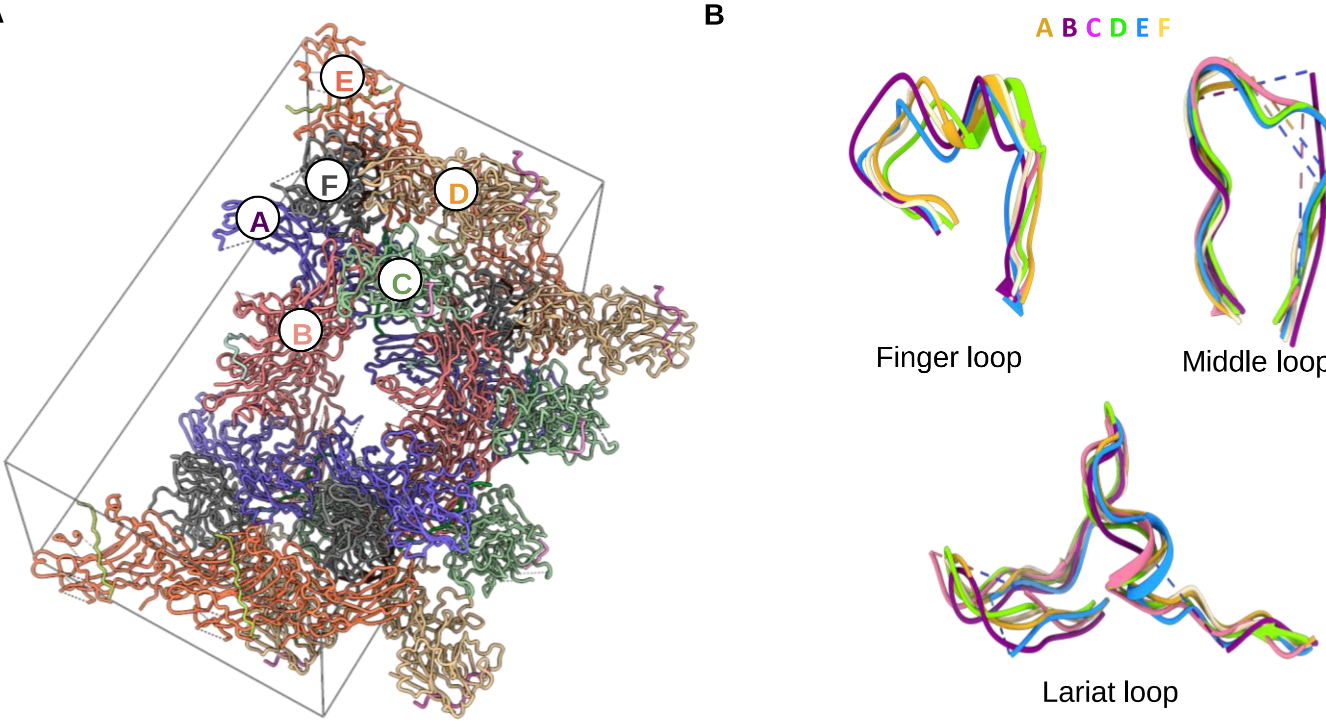


**C**

| βarr2 (Chain A) | CXCR7pp2 (Chain U) |
|---|---|
| R148 | pS350 |
| E146, R166 | pT352 |
| K12, R26, K296 | pS355 |

| βarr2 (Chain B) | CXCR7pp2 (Chain V) |
|---|---|
| R148, R166 | pS350 |
| R166 | pT352 |
| R26 | pS355 |

| βarr2 (Chain C) | CXCR7pp2 (Chain W) |
|---|---|
| H160, R148, K161 | pS350 |
| E67 | pT352 |

| βarr2 (Chain D) | CXCR7pp2 (Chain X) |
|---|---|
| E146, R148 | pS350 |
| K161, R166 | pT352 |
| R166 | pS355 |

| βarr2 (Chain E) | CXCR7pp2 (Chain Y) |
|---|---|
| R148 | pS350 |
| E67 | pT352 |
| K11, K12, R26 | pS355 |

| βarr2 (Chain F) | CXCR7pp2 (Chain Z) |
|---|---|
| K161, R148 | pS350 |
| K161 | pT352 |
| - | pS355 |

**Figure S5. Crystal contact, inter-domain rotation and phosphate interactions in CXCR7-βarr2 structure. A.** Overall arrangement of the six different βarr2 chains in the crystal structure. **B.** Comparison of βarr2 loops in different chains in the asymmetric unit. **C.** The interaction of three phosphate groups in CXCR7pp2 with Lys/Arg residues in different chains of βarr2.

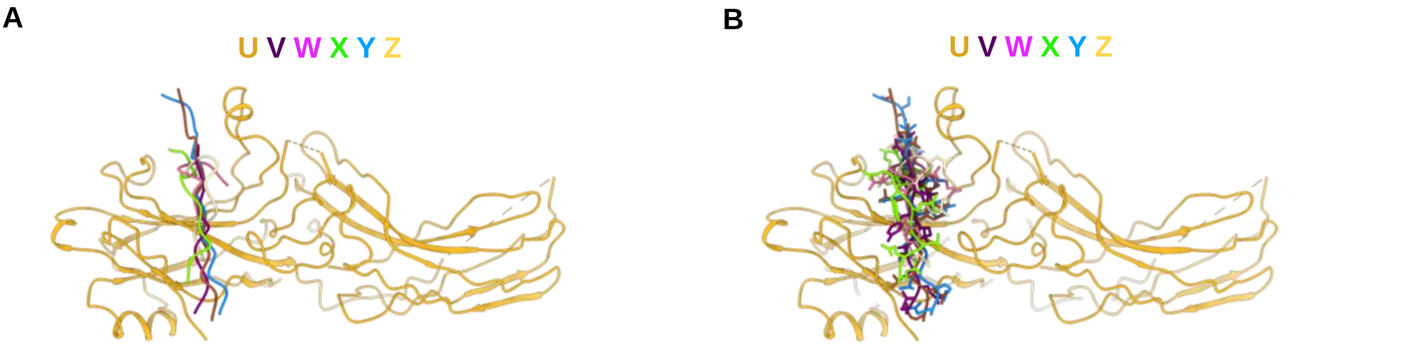


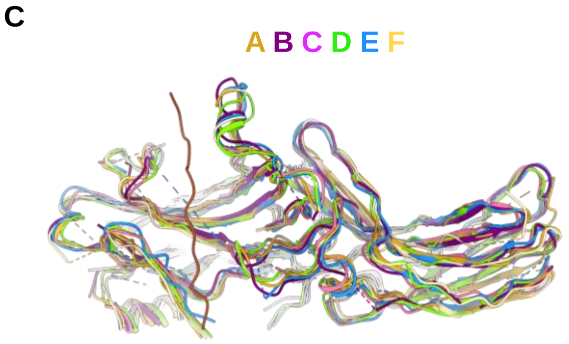


D

| Chains | RMSD (Å) |
|---|---|
| **A-B** | 1.42 |
| **A-C** | 2.991 |
| **A-D** | 1.725 |
| **A-E** | 1.776 |
| **A-F** | 1.145 |

| Chains | RMSD (Å) |
|---|---|
| **U-V** | 1.792 |
| **U-W** | 4.666 |
| **U-X** | 6.345 |
| **U-Y** | 0.819 |
| **U-Z** | 6.097 |

**Figure S6. Comparison of different CXCR7pp2 and βarr2 chains in the asymmetric unit. (A-B) O**verall docking of different chains of CXCR7pp2 with chain D of βarr2 depicted as the template. **C** Overlay of the different chains of βarr2 with chain A of the CXCR7pp2 in the binding pocket. **D.** Relative global RMSD values for the indicated chains of βarr2 and CXCR7pp2 as observed in the crystal structure.

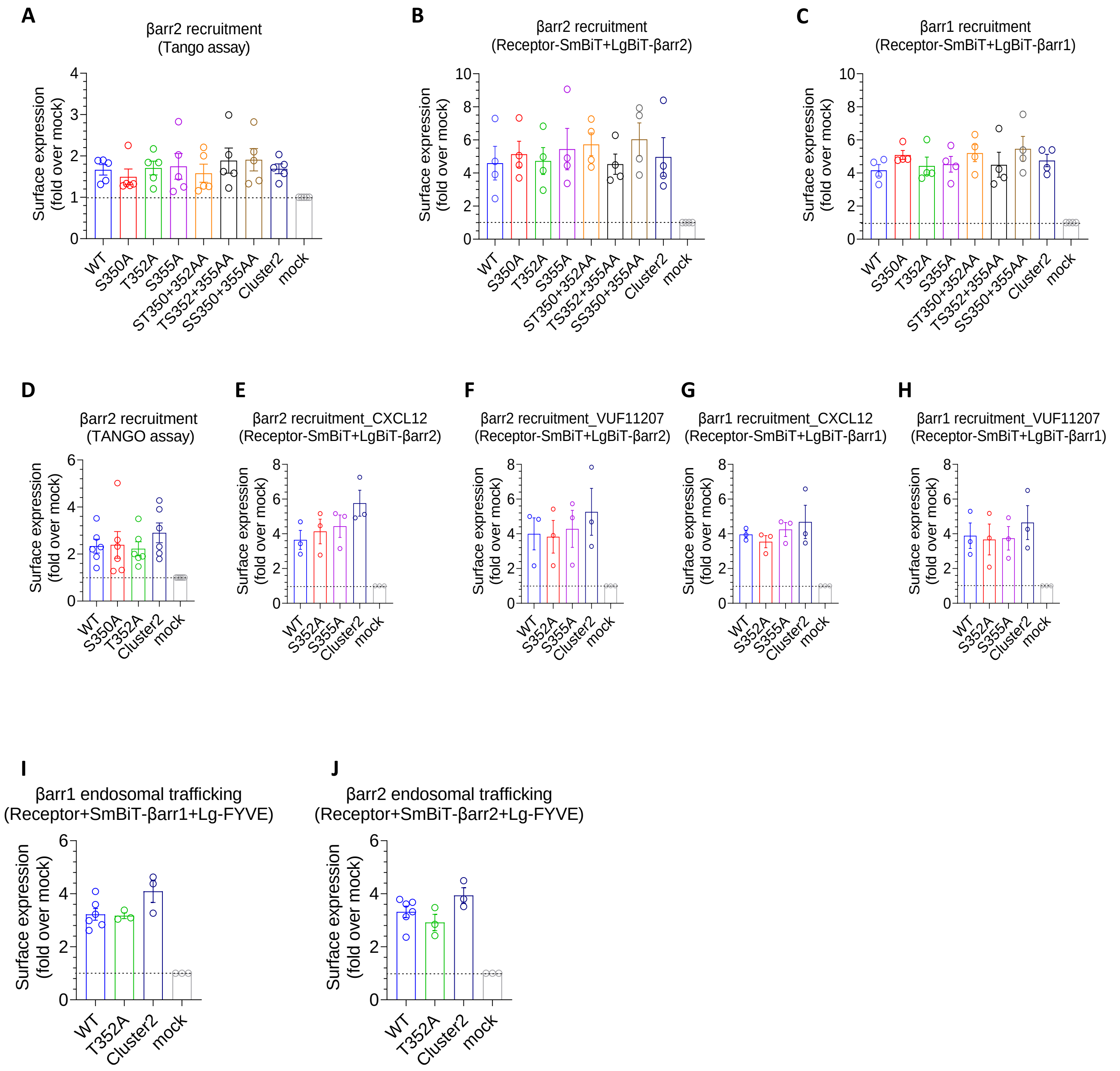


**Figure S7. Surface expression of CXCR7 mutants in different assays. A-E.** Surface expression of indicated receptors were measured in different assays using whole cell ELISA (mean±SEM; n=5 for panel A, n=4 for panel B and C, n=6 for panel D, n=3 for panel E-H and n=3-7 for panel I-J; normalized as fold over mock-transfection).

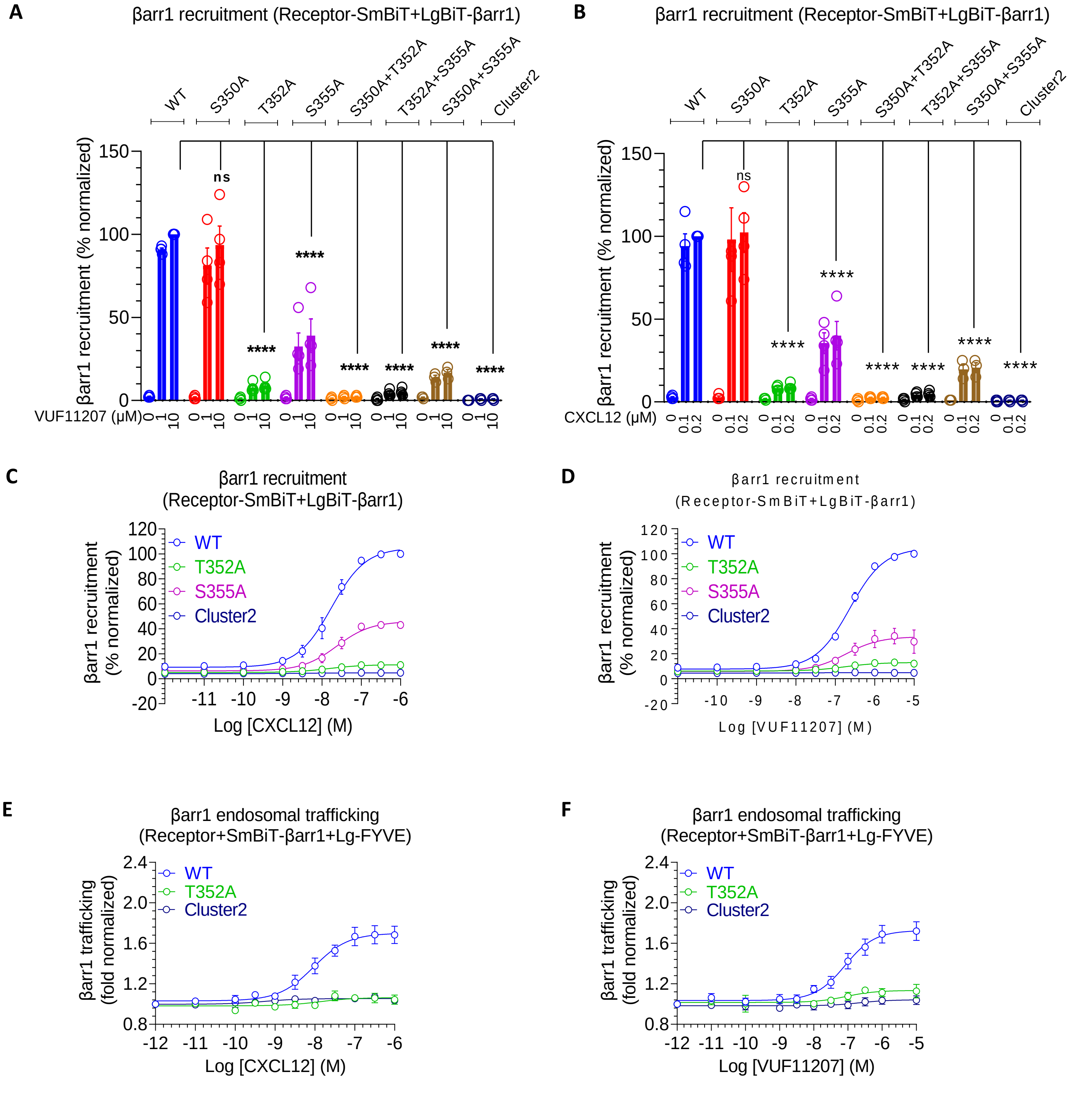


**Figure S8. Contribution of the key phosphorylation sites in CXCR7 on βarr1 recruitment and trafficking. A-B.** CXCL12- and VUF11207-induced βarr1 recruitment, respectively, to the indicated phosphorylation site mutants of CXCR7 using the NanoBiT assay (mean±SEM; n=3-4; normalized with luminescence signal at maximal ligand dose for wild-type as 100%, Two-way ANOVA, Tukey's multiple comparison test, **** p<0.0001, ns= non-significant). **C-D.** Dose response curves of CXCL12- and VUF11207-induced βarr1 recruitment to selected phosphorylation site mutants of CXCR7 in the NanoBiT assay (receptor-SmBiT+LgBiT-βarr1) (mean±SEM; n=3; normalized with luminescence signal at maximal ligand dose as 100%). **E-F.** Dose response curves of CXCL12- and VUF11207-induced βarr1 trafficking to the endosomes for the selected phosphorylation site mutants of CXCR7 in the NanoBiT assay (receptor+SmBiT-βarr1+LgBiT-FYVE) (mean±SEM; n=3-7 for panel E and n=4 for panel F; normalized with the luminescence signal at minimal ligand dose as 1).

**Supplementary Table S1. Statistics for Data Collection and Refinement.**

| Data set | CXCR7pp2-βarr2 |
|---|---|
| ***A. Data collection*** | |
| X-ray source | PLS 7A |
| X-ray wavelength (Å) | 1.0000 |
| Space group | *C*2 |
| Unit cell length (*a*, *b*, *c*, Å) | 91.1 127.2 205.8 |
| Unit cell angle (α, β, γ, °) | 90, 90, 90 |
| Resolution range (Å) | 42.3 – 1.99 |
| Total / unique reflections | 1,172,561 / 313,365 |
| Completeness (%) | 99.6 (98.8)[a] |
| Average $I/\sigma(I)$ | 8.0 (3.6)[a] |
| $R_{merge}$[b] (%) | 12.6 (27.6)[a] |
| ***B. Model refinement statistics*** | |
| Resolution range (Å) | 38.4 - 2.80 |
| $R_{work}$ / $R_{free}$[c] (%) | 28.0 / 33.2 |
| Reflections used in refinement | 57766 |
| Reflections used for R-free | 3006 |
| Average B-factor | 15.2 |
| Number of nonhydrogen atoms | 16131 |
| Macromolecules | 15660 |
| Solvent | 471 |
| Protein residues | 1905 |
| R.m.s. deviations from ideal geometry | |
| Bond lengths (Å) | 0.009 |
| Bond angles (°) | 1.296 |
| Protein-geometry analysis | |
| Ramachandran preferred (%) | 77.8 |
| Ramachandran allowed (%) | 21.1 |
| Ramachandran outliers (%) | 1.1 |

**Footnotes for Table S1**

[a] Values in parentheses refer to the highest resolution shell.
[b] $R_{merge} = \Sigma_{hkl}\Sigma_i \mid I_i(hkl) - \langle I(hkl)\rangle \mid / \Sigma_{hkl}\Sigma_i I_i(hkl)_i$, where $I(hkl)$ is the intensity of reflection *hkl*, $\Sigma_{hkl}$ is the sum over all reflections, and $\Sigma_i$ is the sum over i measurements of reflection *hkl*.
[c] $R = \Sigma_{hkl} \mid |F_{obs}| - |F_{calc}| \mid / \Sigma_{hkl} |F_{obs}|$, where $R_{free}$ was calculated for a randomly chosen 5% of reflections, which were not used for structure refinement and $R_{work}$ was calculated for the remaining.
